# The *CYP71A*, *NIT*, *AMI,* and *IAMH* gene families are dispensable for indole-3-acetaldoxime-mediated auxin biosynthesis in Arabidopsis

**DOI:** 10.1101/2025.06.03.657619

**Authors:** M. Fenech, J. Brumos, A. Pěnčík, B. Edwards, S. Belcapo, J. DeLacey, A. Patel, M.M. Kater, X. Li, K. Ljung, O. Novak, J.M. Alonso, A.N. Stepanova

**Affiliations:** Department of Plant and Microbial Biology, College of Agriculture and Life Sciences, North Carolina State University, Raleigh, North Carolina, United States of America; Institute for Plant Molecular and Cell Biology (IBMCP), CSIC-Universitat Politècnica de Valencia, Valencia, Spain; Laboratory of Growth Regulators, Institute of Experimental Botany, The Czech Academy of Sciences & Faculty of Science, Palacký University, Olomouc, Czech Republic; Department of BioScience, Università degli Studi di Milano, Milan, Italy; Department of Forest Genetics and Plant Physiology, Umeå Plant Science Centre (UPSC), Swedish University of Agricultural Sciences, Umeå, Sweden

**Author notes:** **Corresponding authors**, Alonso J.M. and Stepanova A.N. The authors responsible for distribution of materials integral to the findings presented in this article in accordance with the policy described in the Instructions for Authors (https://academic.oup.com/plcell/pages/General-Instructions) are: Anna N. Stepanova and Jose M. Alonso.

## Abstract

Indole-3-acetic acid (IAA) is a crucial auxin governing plant development and environmental responses. While the indole-3-pyruvic acid (IPyA) pathway is the predominant IAA biosynthesis route, other pathways, like the indole-3-acetaldoxime (IAOx) pathway, have been proposed. The IAOx pathway has garnered attention due to its supposed activation in auxin-overproducing mutants (e.g., *sur1, sur2, ugt74b1*) and the auxin-like responses triggered by exogenous application of its proposed intermediates: IAOx, indole-3-acetonitrile (IAN), and indole-3-acetamide (IAM). However, despite supporting evidence for individual steps, conclusive physiological relevance of the IAOx pathway remains unproven. Using a comprehensive genetic approach combined with metabolic and phenotypic profiling, we demonstrate that mutating gene families proposed to function in the IAOx pathway does not result in prominent auxin-deficient phenotypes, nor are these genes required for high-auxin production in the *sur2* mutant. Our findings also challenge the previously postulated linear IAOx pathway. While exogenously provided IAOx, IAN, and IAM can be converted to IAA *in vivo*, they do not act as precursors for each other. Finally, our findings question the physiological relevance of IAM and IAN as IAA precursors in plants and suggest the existence of a yet uncharacterized auxin biosynthetic route, likely involving IAOx as an intermediate, for the production of IAA in the *sur2* mutant. Future identification of the metabolic steps and the corresponding genes in this new pathway may uncover the previously unknown way of synthesizing IAA in plants.

## Introduction

The plant hormone auxin plays a central regulatory role in almost every aspect of the plant life cycle, from activating developmental programs to triggering responses to environmental cues (Zhao, 2018). The best-characterized auxin, indole-3-acetic acid (IAA), is known to act in a concentration-dependent manner, and thus, its abundance is spatiotemporally regulated through finely controlled and interconnected biosynthetic, transport, and degradation processes (Petricka et al., 2012; Brumos et al., 2018; Hayashi et al., 2021; Zhang et al., 2023). Different genetic, biochemical, metabolic, and phenotypic approaches have been used to identify and characterize the key elements in these processes (Vorwerk et al., 2001; Pollmann et al., 2002; Pollmann et al., 2006; Müller et al., 2015). In the case of auxin biosynthesis, two general routes of IAA production have been proposed based on whether or not they use the amino acid tryptophan (TRP) as a precursor (Figure 1). The tentative TRP-independent pathway was originally proposed based on radiolabeled indole feeding experiments in maize and Arabidopsis TRP biosynthetic mutants (Wright et al., 1991; Normanly et al., 1993; Ostin et al., 1999; Ouyang et al., 2000). These experiments suggested that IAA could be produced not from TRP but from indole derived from upstream TRP precursors. Despite considerable efforts, the identification of the enzymatic activities and the corresponding genes involved in this pathway has proven elusive. Thus, although a cytosolic indole synthase potentially involved in the conversion of indole-3-glycerol phosphate (IGP) into indole has been identified (Li et al., 1995; Ouyang et al., 2000; Wang et al., 2015), the mechanisms by which indole would then be converted into IAA and the overall relevance of this pathway remain controversial (Nonhebel, 2015). In addition to the disputed TRP-independent pathway, several somewhat interconnected TRP-dependent routes for IAA production in plants have been proposed (Figure 1), each with different levels of experimental support and scientific community acceptance (Nonhebel, 2015). These TRP-dependent pathways are typically referred to by the name of a key biosynthetic intermediate.

**Figure 1.**
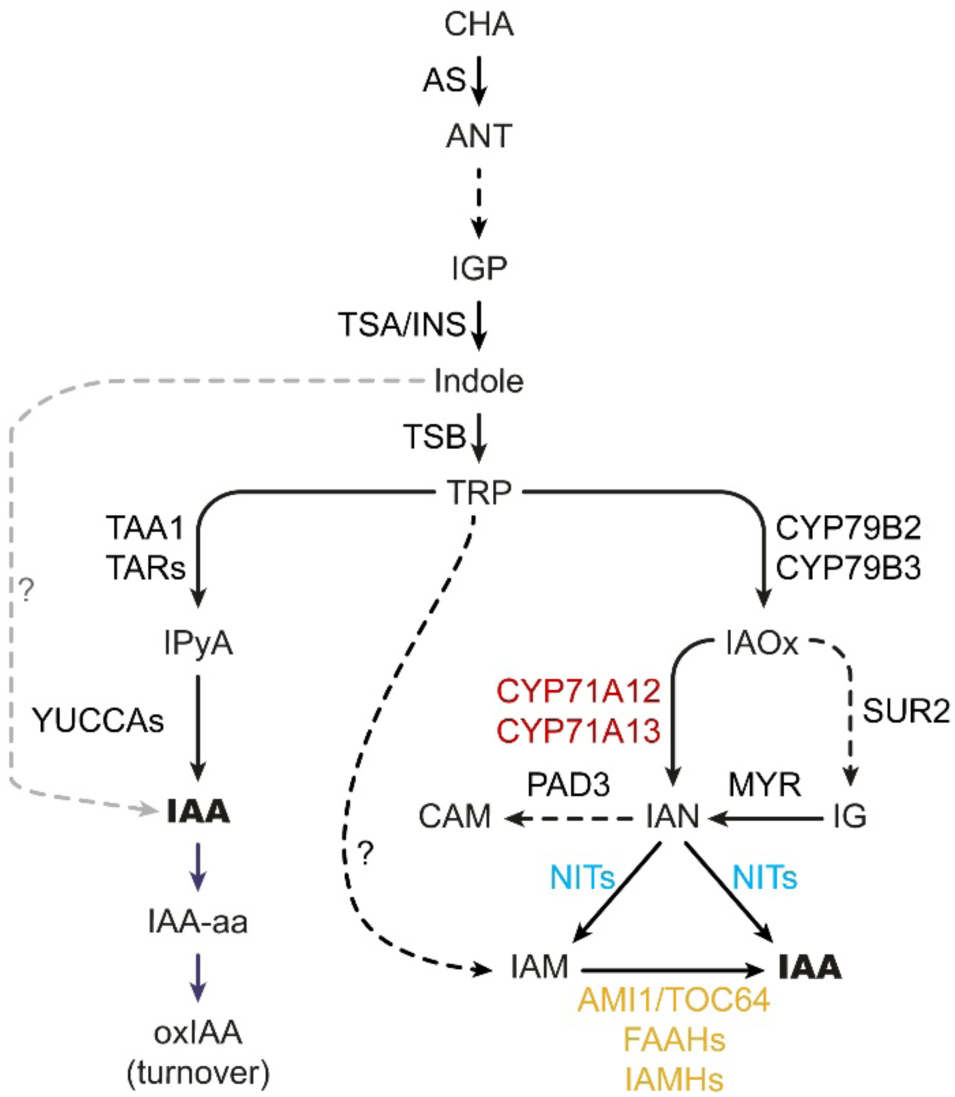
A model of the auxin biosynthesis pathway leading to the production of indole-acetic acid (IAA). Tryptophan (TRP)-dependent and TRP-independent routes are marked with black and gray arrows, respectively. Solid arrows indicate single-step conversions, while dashed arrows represent multi-step conversions. TRP-dependent indole-pyruvic acid (IPyA) and putative indole acetaldoxime (IAOx) routes are depicted, as are the abbreviated versions of the upstream TRP biosynthesis out of chorismic acid (CHA) and the competing parallel indole glucosinolate (IG) and camalexin (CAM) biosynthesis routes that split at IAOx and indole-3-acetonitrile (IAN), respectively. The CYP71A, NIT and AMI1/TOC64/FAAH protein families examined in this work as putative components of the IAOx pathway are displayed in red, blue, and yellow, respectively. CHA: chorismic acid, ANT: anthranilate, IGP: indole-3-glycerol phosphate, IAA-aa: amino-acid-conjugated IAA (e.g., indole-3-acetyl-aspartate (IAA-Asp) and indole-3-acetyl-glutamate (IAA-Glu)), oxIAA: 2-oxindole-3-acetic acid, IAM: indole-3-acetamide; AS: anthranilate synthase, TSA: TRP SYNTHASE α (chloroplast), INS: INDOLE SYNTHASE (cytosol), TSB: TRP SYNTHASE β, TAA1: TRP AMINOTRANSFERASE OF ARABIDOPSIS1 (WEI8), TAR: TAA1-RELATED (TAR1, TAR2), NIT: NITRILASE (NIT1-NIT4), AMI1: AMIDASE1 (TOC64-I), TOC64: TRANSLOCON OF OUTER MEMBRANE OF THE CHLOROPLAST64 (*TOC64-III*, *TOC64-V*), FAAH: FATTY ACID AMIDE HYDROLASE (FAAH1-FAAH4), IAMH: IAM HYDROLASE (IAMH1, IAMH2), MYR: MYROSYNASE, PAD3: PHYTOALEXIN DEFICIENT3, ?: unknown enzyme(s).

The best characterized of these TRP-dependent routes of auxin production is the indole-3-pyruvic acid (IPyA) pathway that involves the conversion of TRP into IPyA by a small family of aminotransferases (TRP AMINOTRANSFERASE OF ARABIDOSPIS1 (TAA1) and TAA1-RELATED (TARs)) and IPyA into IAA by a family of flavin-containing monooxygenases (YUCCAs). Evidence in support of this pathway is, by far, the most conclusive and includes genetic, metabolic, and phylogenetic results that together substantiate the universality and preponderance of this route among plants (Zhao et al., 2001; Stepanova et al., 2008; Tao et al., 2008; Yamada et al., 2009; Stepanova et al., 2011; Zhou et al., 2011; Eklund et al., 2015). In contrast, the experimental support for the rest of TRP-dependent pathways is somewhat fragmentary and inconclusive. For instance, although radiolabeled precursor feeding experiments indicate that key components of these other pathways—such as indole-3-acetaldoxime (IAOx), indole-3-acetonitrile (IAN), indole-3-acetamide (IAM), tryptamine (TAM), and indole-3-acetaldheyde (IAAld)—can all serve as intermediates in the production of IAA from TRP (Seo et al., 1998; Pollmann et al., 2002; Pollmann et al., 2006; Sugawara et al., 2009; Lehmann et al., 2010; Novák et al., 2012), conclusive identification of the enzymatic reactions and genes involved, and importantly, evidence supporting the physiological relevance of these pathways, are rather limited.

Support for the role of IAN as a TRP-derived IAA precursor comes not only from the radiolabeling experiments mentioned above, but also from experimental evidence that a number of phylogenetically distant species have the ability to convert exogenous IAN into IAA (Thimann and Mahadevan, 1964). Endogenous IAN has been reported for species belonging to the *Brassicacea* family, like cabbage or Arabidopsis (Henbest et al., 1953; Rajagopal and Larsen, 1972; Sugawara et al., 2009), but also in other species belonging to *Amaranthaceae*, *Asteraceae/Compositae*, *Convolvulaceae*, *Cucurbitaceae*, *Lamiaceae/Labiatae*, *Fabaceae/Leguminosae*, *Araceae/Lemnaceae*, or *Liliaceae* families (Rajagopal and Larsen, 1972), and in coleoptiles of monocots (*Poaceae/Gramineae*) like maize and oat (Rajagopal and Larsen, 1972; Park et al., 2003). The identification of a set of Arabidopsis mutants impaired in their response to exogenous IAN shed more light on the potential role of IAN as an auxin precursor (Normanly et al., 1997). Characterization of these mutants identified *NITRILASE1* (*NIT1*) as a gene involved in the plant response to exogenous IAN. *NIT1* is a member of a small family of four genes in Arabidopsis, *NIT1* through *NIT4*, of which *NIT1 to NIT3* are thought to function in IAN metabolism, whereas *NIT4* has been implicated in cyanide detoxification (Vorwerk et al., 2001; Mulelu et al., 2019). Although NIT1 to NIT3 can convert IAN into IAA *in vitro*, they show much greater affinity and catalytic rate transforming other naturally occurring substrates, like 3-phenylpropionitrile, allylcyanide, (phenylthio)acetonitrile, or (methylthio)acetonitrile than they do for IAN (Vorwerk et al., 2001). Importantly, neither the loss-of-function of *NIT1* nor the knockdown RNAi lines for *NIT1* to *NIT3* showed obvious phenotypic defects beyond the resistance to exogenous IAN (Normanly et al., 1997; Lehmann et al., 2017), casting some doubts about the role of endogenous IAN as an IAA precursor and the potential physiological relevance of a native IAN-dependent IAA biosynthetic pathway. Nevertheless, due to functional redundancy among the *NIT* gene family members and the potential residual gene activity in the RNAi knockdowns, the lack of obvious phenotypes of these mutants has not formally ruled out a potential role of this gene family in the production of IAA during normal plant development.

As with IAN, a variety of plant species have been shown to contain IAM (Pollmann et al., 2003; Pollmann et al., 2006; Lehmann et al., 2010) and to have the capability of uptaking exogenous IAM and converting it into IAA (Pollmann et al., 2003; Pollmann et al., 2006; Lehmann et al., 2010). The isolation and characterization of AMIDASE1 (AMI1), an Arabidopsis amidase capable of hydrolyzing IAM *in vitro* (Pollmann et al., 2003; Pollmann et al., 2006), along with its paralogs from the most-closely related *TRANSLOCON OF THE OUTER MEMBRANE OF THE CHLOROPLASTS64* (*TOC64*) gene family (*TOC64-III* and *TOC64-V*; (Pollmann et al., 2006; Aronsson et al., 2007) and a more distantly related *FATTY ACID AMIDE HYDROLASE* (*FAAH*) family (Pollmann et al., 2006; Keereetaweep et al., 2013), provided an opportunity for testing the potential role of the IAM pathway in plants. As in the case of the *NIT*s, the lack of a systematic mutant analysis of *AMI1* and related genes has precluded the conclusive placement of these genes in a hypothetical IAM route of auxin biosynthesis. More recently, a new *IAM HYDROLASE* (*IAMH1*), unrelated in sequence to *AMI1*, was identified in Arabidopsis using genetic screening for IAM-resistant mutants and implicated in the conversion of IAM to IAA (Gao et al., 2020). *IAMH1* has a close paralog in Arabidopsis, *IAMH2*, that, based on double mutant analysis, is also involved in the production of IAA in plants treated with IAM. Importantly, although these double mutant plants show clear resistance to exogenous IAM, they do not display prominent auxin-related developmental defects under normal laboratory growth conditions (Gao et al., 2020), questioning their role in the proposed IAM-dependent IAA biosynthetic pathway.

In contrast with IAM, IAOx has only been detected in some plant species belonging to the *Brassicaceae* family (Cooney and Nonhebel, 1991; Sugawara et al., 2009) and in two species of the genus *Erythroxylum* (*Erythroxylaceae*; Luck et al., 2016). Consistent with this, the presence of close homologs of the Arabidopsis *CYP79B* gene family involved in the conversion of TRP to IAOx has also been found only in the Brassicales order (Bak et al., 2001). In these plant species, IAOx is a precursor in the production of two defense compounds: indole glucosinolates (IG; Bak et al., 1998; Hull et al., 2000; Mikkelsen et al., 2000; Bak et al., 2001; Zhao et al., 2002) and camalexins (CAM; Glawischnig et al., 2004; Nafisi et al., 2007; Müller et al., 2015). Importantly, several enzymatic reactions and the corresponding genes involved in these two pathways have been identified and characterized in Arabidopsis. Thus, the first step in the indole glucosinolate pathway consists of the conversion of TRP into IAOx by two partially redundant cytochrome P450s, CYP79B2 and CYP79B3 (Hull et al., 2000; Mikkelsen et al., 2000; Zhao et al., 2002), a step that is shared with the CAM biosynthesis pathway (Glawischnig et al., 2004). Accordingly, the double mutant *cyp79b2 cyp79b3* completely lacks these two important defense compounds (Glawischnig et al., 2004). IAOx can then be channeled into the production of CAM by a small family of cytochrome P450s related to CYP71A13 (Nafisi et al., 2007; Müller et al., 2015) or into IG by CYP83B1 (SUPERROOT2, SUR2) (Bak et al., 2001). Like the conversion of TRP into IAOx by the CYP79B2/3, the mutant analysis provides strong support for the involvement of SUR2 and CYP71A12/13 in the first committed steps of IG and CAM biosynthesis, respectively, as the production of these defense compounds was greatly reduced in the corresponding mutants (Zhao et al., 2002; Nafisi et al., 2007; Müller et al., 2015).

The connection of CAM and IAA biosynthesis pathways stems from the involvement of IAN as the product of IAOx dehydration catalyzed by CYP71A12/13 (Mucha et al., 2019). On the other hand, the link between IG and IAA biosynthesis comes from the high-auxin phenotypes observed not only in the *sur2* mutant but also in several of the mutants affecting downstream IG pathway genes such as *SUR1* (Boerjan et al., 1995) and *UDP-GLUCOSYLTRANSFERASE 74B1* (*UGT74B1)* (Grubb et al., 2004). Furthermore, feeding experiments using radiolabeled IAOx in the *cyp79b2 cyp79b3* mutant identified IAN and IAM as likely conversion products of IAOx (Sugawara et al., 2009). These results, together with the current understanding of the role of *CYP71As*, *NITs*, *AMI* and *IAMHs* in the metabolism of IAOx, IAN, and IAM, respectively, have been used to propose a metabolic and genetic pathway for the production of IAA from IAOx (Sugawara et al., 2009). In this pathway, IAOx would be first converted into IAN by the CYP71As, NITs would then catalyze the conversion of IAN into IAM and/or IAA, and finally, AMI and/or IAMH would catalyze the formation of IAA from IAM (Figure 1).

The IAOx pathway has attracted considerable attention due to the fact that its activation in *sur2* and other IG mutants results in plants with high IAA levels and strong auxin overproduction phenotypes (Boerjan et al., 1995; Seo et al., 1998; Barlier et al., 2000; Grubb et al., 2004) suggesting a potential physiologically relevant alternative to the IPyA pathway of IAA production. On the other hand, the general importance of the IAOx route of auxin production is somewhat diminished by its restriction to mostly plant species of the *Brassicaceae* family and the lack of prominent developmental defects in the *cyp79b2 cyp79b3* double knockout mutants where this pathway is completely blocked (Stepanova et al., 2011; Tsugafune et al., 2017). It is important to emphasize, however, that plants as distantly related to Arabidopsis as maize have been shown to contain the enzymatic machinery necessary to convert exogenous IAOx into IAA (Perez et al., 2021), and *Medicago truncatula* plants treated with IAOx show strong high-auxin phenotypic responses (Buezo et al., 2019; Roman et al., 2023). Furthermore, although no homologous sequence to *CYP79B2* or *CYP79B3* can be found in the maize genome, cytochrome P450s of the CYP79A family have been shown to be able to produce IAOx and phenyl-acetaldoxime (PAOx) in maize and sorghum, respectively (Irmisch et al., 2015; Perez et al., 2023), leaving open the possibility for a functional IAOx-related pathway in a broader range of plant species.

Thus, the current view of the IAA production pathways that may function in plants in parallel with the primary IPyA-dependent route is confusing, with different levels of support and undermining experimental evidence having been put forward. One key limitation to addressing this problem has been the lack of mutant lines where the whole family of genes putatively involved in these alternative IAA production pathways has been knocked out. Thus, as the first step towards clarifying the role of the different IAA biosynthetic pathways, we generated high-order mutant lines for each of three key gene families, *CYP71A12/13/18, NIT1/2/3/4*, and *AMI1*/*TOC64*/*FAAH* that in combination with previously characterized mutants (such as *sur2, cyp79b2 cyp79b3,* and *iamh1 iamh2*) and a comprehensive set of phenotypic and metabolic approaches has allowed us to thoroughly test the role of these gene families in auxin biosynthesis. Our results clearly indicate that none of the three gene families investigated here play a role in the IAOx pathway activated in the *sur2* mutant and question the validity of the previously proposed linear pathway for the conversion of IAOx into IAA via IAN and IAM. We anticipate this work will trigger new efforts to unveil how excess auxin is produced in *sur2* and investigate the possible physiological relevance of such a pathway in *Brassicaceae* and beyond.

## Results

### Generation of mutant lines

In addition to the predominant IPyA auxin biosynthetic pathway, a second TRP-dependent route to produce IAA via IAOx, IAN, and IAM has been proposed in Arabidopsis and other *Brassicaceae* species (Figure 1) (Sugawara et al., 2009). The enzymes thought to catalyze the corresponding reactions are encoded by three multigenic families represented in Arabidopsis by the *CYP71A*s (Nafisi et al., 2007; Müller et al., 2015), *NIT*s (Normanly et al., 1997), and *AMI*s (Lehmann et al., 2010). Although several lines of evidence support individual steps of this metabolic pathway as described above, the possible connections between the individual steps and the overall functionality of this pathway have not been thoroughly investigated. Thus, in order to test the potential contribution of the proposed IAOx-dependent pathway to the IAA pools in Arabidopsis, we decided to take a genetic approach and examine the phenotypic and metabolic consequences of knocking out the three enzymatic steps mentioned above. Due to potential functional redundancy, we first identified the closest family members of the three better-characterized enzymes in the IAOx pathway: *CYP71A13*, *NIT1*, and *AMI1*. Amino acid sequence comparison and phylogenetic analysis (Supplementary Fig. S1) led us to identify three genes as part of the CYP71A family (*A12*, *A13*, and *A18*), four members in the *NIT* family (*NIT1-NIT4*), and, consistent with previous reports, seven extended *AMI* family members that include three *TOC64*s and four *FAAH*s (Pollmann et al., 2006) potentially involved in IAOx-dependent auxin biosynthesis.

To test the involvement of these three gene families in IAA production, we used a combination of publicly available T-DNA lines and targeted genomic editing tools to generate the corresponding high-order gene family mutants (Supplementary Fig. S2; Supplementary Fig. S3). We observed embryo lethality in *faah3* single mutants (Supplementary Fig. S4) and therefore, the highest order mutant for the *AMI* gene family was propagated as a segregating line, *ami1-1 toc64-III toc64-V faah1 faah2 faah3/+ faah4*. The knockouts for the whole *CYP71A* and *NIT* families were fully viable and fertile. Considering the strong phenotype described for the IPyA-deficient *TAA1/TAR* family double mutant *wei8 tar2* and the early development arrest of *wei8 tar1 tar2* triple mutant (Stepanova et al., 2008), we hypothesized that the contribution of the IAOx pathway to the auxin pools in plants grown under standard laboratory conditions would be, at best, minor. Therefore, to enhance any potential auxin-related phenotypes that plants lacking functional *CYP71A*, *NIT*, and *AMI1/TOC64/FAAH* gene families may have, we genetically activated the metabolic flux of the proposed IAOx pathway by introgressing the *sur2* T-DNA allele (Stepanova et al., 2005) into the three high-order gene family knockout lines mentioned above. Due to difficulties in combining the existing mutant alleles of *faah4* and *sur2* because of genetic linkage, the fairly distant phylogenetic relationship between the founding family member AMI1 and FAAH4 (Supplementary Fig. S1), and the embryo lethality of *faah3* (Supplementary Fig. S4), the highest order mutant for the *AMI* family in the *sur2* background we built was *ami1-2 toc64-III toc64-V faah1 faah2 sur2*. Then, to determine the potential involvement of these three gene families in auxin production in the wild type (WT), as well as in the *sur2* mutant background, we carried out a battery of phenotypic and metabolomic analyses of the higher order mutants of each gene family with and without *sur2*. We reasoned that the role of individual genes could be dissected at a later stage once a specific phenotype was observed for the high-order mutants, hereon, referred to as IAOx high-order mutants.

### Mutants with defects in the postulated IAOx pathway do not show auxin-related defects and fail to suppress high-auxin phenotypes of *sur2*

To determine the potential involvement of *CYP71A*, *NIT*, and *AMI1/TOC64/FAAH* gene families in auxin biosynthesis, we examined a battery of auxin-related phenotypes of their high-order mutants using the mild auxin-deficient mutant *wei8-1* (Stepanova et al., 2008; Tao et al., 2008; Yamada et al., 2009) as a positive control for subtle auxin-related defects (Figure 2A; Supplementary Fig. S5A-D). The mild auxin-deficient mutant *wei8* did not display any major developmental defects in seedling organ size (hypocotyl and root length) when grown in constant darkness (3 days; Supplementary Fig. S5A, C) or under continuous light (7 days; Supplementary Fig. S5B, D). However, *wei8* roots were significantly shorter than those of WT when three-day-old dark-grown seedlings were transferred to continuous light for four additional days (Figure 2A, B). In contrast, none of the high-order IAOx mutants showed significant differences in seedling organ size compared to WT under any of the growth conditions tested (Figure 2A-B; Supplementary Fig. S5A-D). Furthermore, neither *wei8* nor IAOx mutants displayed any prominent phenotypic defects in soil-grown adults (Figure 2C).

**Figure 2.**
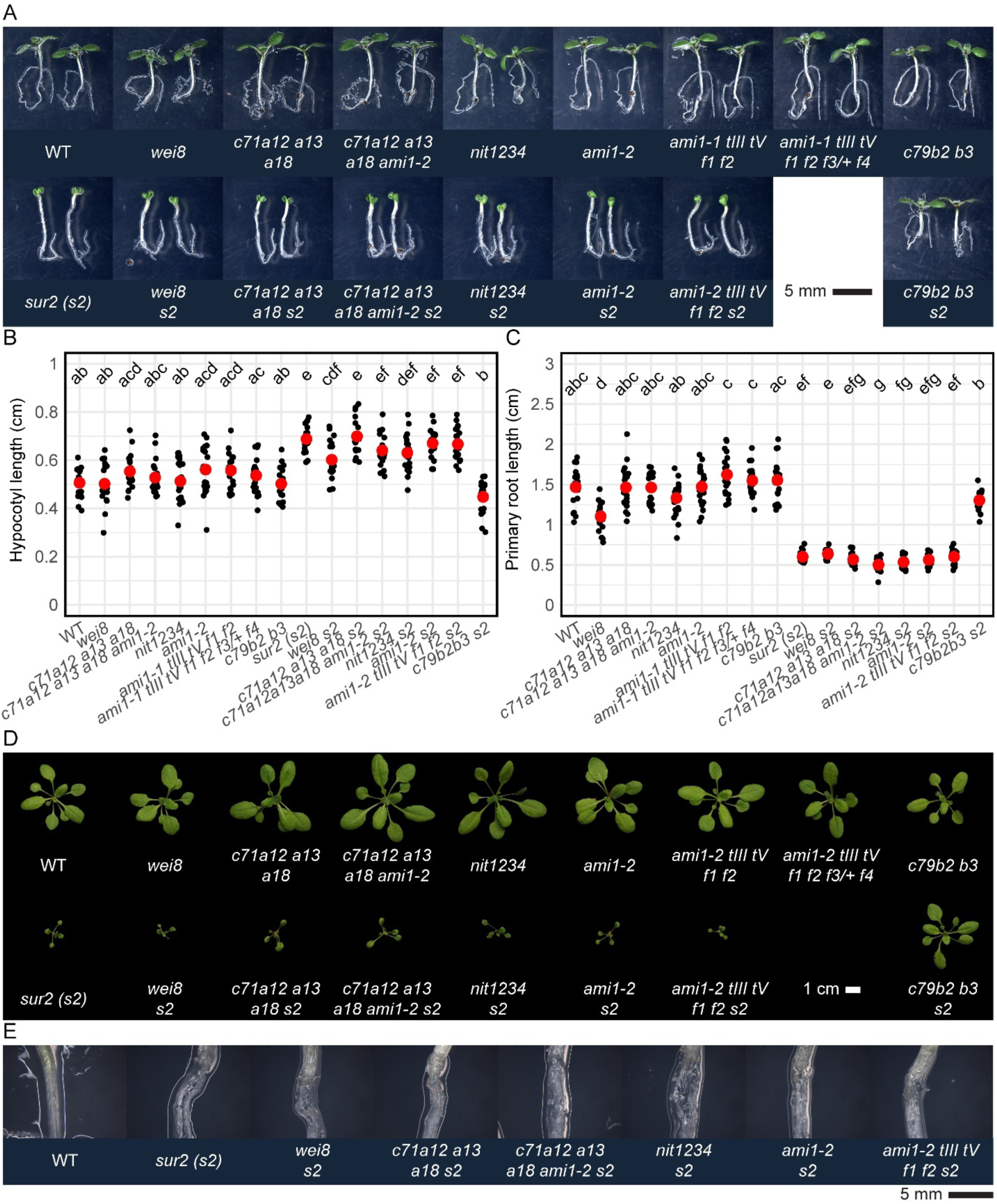
Mutants defective in the proposed IAOx pathway of auxin biosynthesis display no prominent growth defects. (A) Phenotypes of seedlings germinated on horizontal plates in the dark for three days and transferred to continuous light for four additional days. (B) Organ size quantification and statistical analysis of seedlings shown in (A). (C) Rosettes of three-week-old plants grown in soil under long-day conditions (16-h fluorescent light/8-h darkness) after germination on vertical plates for three days in the dark and four days under continuous LED light. (D) The characteristic high-auxin hypocotyl disintegration phenotype of seven-day-old *sur2* seedlings grown for three days in the dark followed by four days in continuous light is not prevented by mutations in genes implicated in the putative IAOx pathway. WT: wild-type (Col-0), *c71a12 a13 a18: cyp71a12 cyp71a13 cyp71a18, c71a12 a13 a18 ami1-2: cyp71a12 cyp71a13 cyp71a18 ami1-2, nit1234: nit1 nit2 nit3 nit4, ami1-1 tIII tV f1 f2: ami1-1 toc64-III toc64-V faah1 faah2, ami1-1 tIII tV f1 f2 f3/+ f4: ami1-1 toc64-III toc64-V faah1 faah2 faah3/+ faah4, s2: sur2, c79b2 b3: cyp79b2 cyp79b3*.

Given the absence of observable defects in high-order IAOx mutants, including the previously characterized *cyp79b2 cyp79b3* known to block the production of IAOx from TRP (Stepanova et al., 2011) (Figure 2), we considered the possibility that the IAOx pathway may not be active at the developmental stages or under the environmental conditions examined. Therefore, we characterized the phenotypes of these mutants when the putative IAOx pathway is genetically activated by knocking out the *SUR2* gene. In this mutant, the flow of IAOx into IG production is blocked, with IAOx now channeled into IAA production (Figure 1) (Barlier et al., 2000; Bak et al., 2001). Overproduction of auxin in *sur2* translated into statistically shorter hypocotyls and roots in three-day-old dark-grown seedlings (Supplementary Fig. S5 A, C) or longer hypocotyls and shorter roots in seven-day-old plants grown in constant light (Supplementary Fig. S5B, D) or in the dark for three days followed by four days in the light (Figure 2A-B). Additionally, high levels of auxin in *sur2* plants led to the development of characteristic small, epinastic cotyledons (Figure 2A) and the disintegration of hypocotyls upon adventitious root emergence in seven-day-old seedlings that underwent a dark-to-light transition (Figure 2D), as well as small, epinastic rosette leaves in soil-grown adults (Figure 2C). As expected, *wei8* did not suppress any of the high-auxin phenotypes of *sur2* (Figure 2; Supplementary Fig. S5) since *WEI8* and *SUR2* are thought to take part in two independent auxin biosynthesis routes. On the other hand, we reasoned that if indeed the IAOx gene families studied in this work were involved in the proposed IAOx-dependent *sur2*-activated auxin biosynthesis pathway, knocking them out should result in the suppression of the *sur2* high-auxin phenotype, as observed in the *cyp79b2/b3 sur2* triple mutant in all the growth conditions tested (Figure 2; Supplementary Fig. S5) (Stepanova et al., 2011). However, none of the IAOx family mutants suppressed any of the *sur2* phenotypes examined above, neither at the seedling stage (Figure 2A-B; Supplementary Fig. S5) nor at the three-week-old rosette stage (Figure 2C).

We looked into the possibility that the T-DNA intronic mutant alleles could be spliced out, with the functionality of the gene fully restored. We found that the expression of *AMI1* in *ami1-1 toc64-III toc64-V faah1 faah2 faah3/+ faah4* mutant was reduced more than 330 fold compared to WT, whereas the levels of *FAAH1* and *FAAH4* were mildly reduced (Supplementary Fig. S6). As all other *CYP71A*, *NIT*, and *AMI1/TOC64* mutants leveraged in this work are exonic knockouts or deletions (Supplementary Fig. S2), in the absence of prominent phenotypes, we conclude that (1) the most closely related members of these gene families examined herein do not play a prominent role in auxin biosynthesis under the studied growth conditions, and (2) their functions are not required for the auxin production activated in *sur2* mutant plants.

### The currently accepted metabolic pathway for the conversion of IAOx to IAA needs to be reassessed

The inability of the high-order IAOx mutants to suppress the *sur2* high-auxin phenotypes suggests that the corresponding enzymes may not be involved in the sequential conversion of IAOx into IAN, IAM, and finally, IAA (Figure 1). Alternatively, these protein families may catalyze the proposed enzymatic reactions, but the metabolic pathway by which excess auxin is produced in *sur2* is different from the previously postulated IAOx-IAN-IAM-IAA route (Sugawara et al., 2009). To clarify this situation, we decided to examine the effects of feeding various auxin precursors and the ability of the different high-order IAOx mutants to block the potential phenotypic consequences of the conversion of these auxin precursors into IAA *in planta*.

To determine the experimental conditions where these compounds trigger clear auxin-like responses, we first examined the auxin-related phenotypes of increasing concentrations of each of the proposed IAA precursors of the IAOx pathway in WT plants in three-day-old dark-grown (Supplementary Fig. S7) and five-day-old light-grown plants (Figure 3; Supplementary Fig. S7). We observed that in both experimental setups, exogenous IAOx and IAN, much like with IAA treatments, led to prominent dose-dependent root growth inhibition in WT plants, whereas IAM treatment led to profound hypocotyl growth promotion specifically in the light (Figure 3; Supplementary Fig. S7). In three-day-old dark-grown WT seedlings exposed to high IAM concentrations of 20 µM and above, mildly shorter root and hypocotyl lengths were observed relative to plants exposed to the solvent (DMSO) (Supplementary Fig. S7). In contrast, in continuous light, 10-60 µM IAM stimulated pronounced (1.5-3-fold) hypocotyl elongation in five-day-old WT seedlings, whereas the root lengths of these plants were inhibited at 20 µM IAM and above by up to 25% (Figure 3).

**Figure 3.**
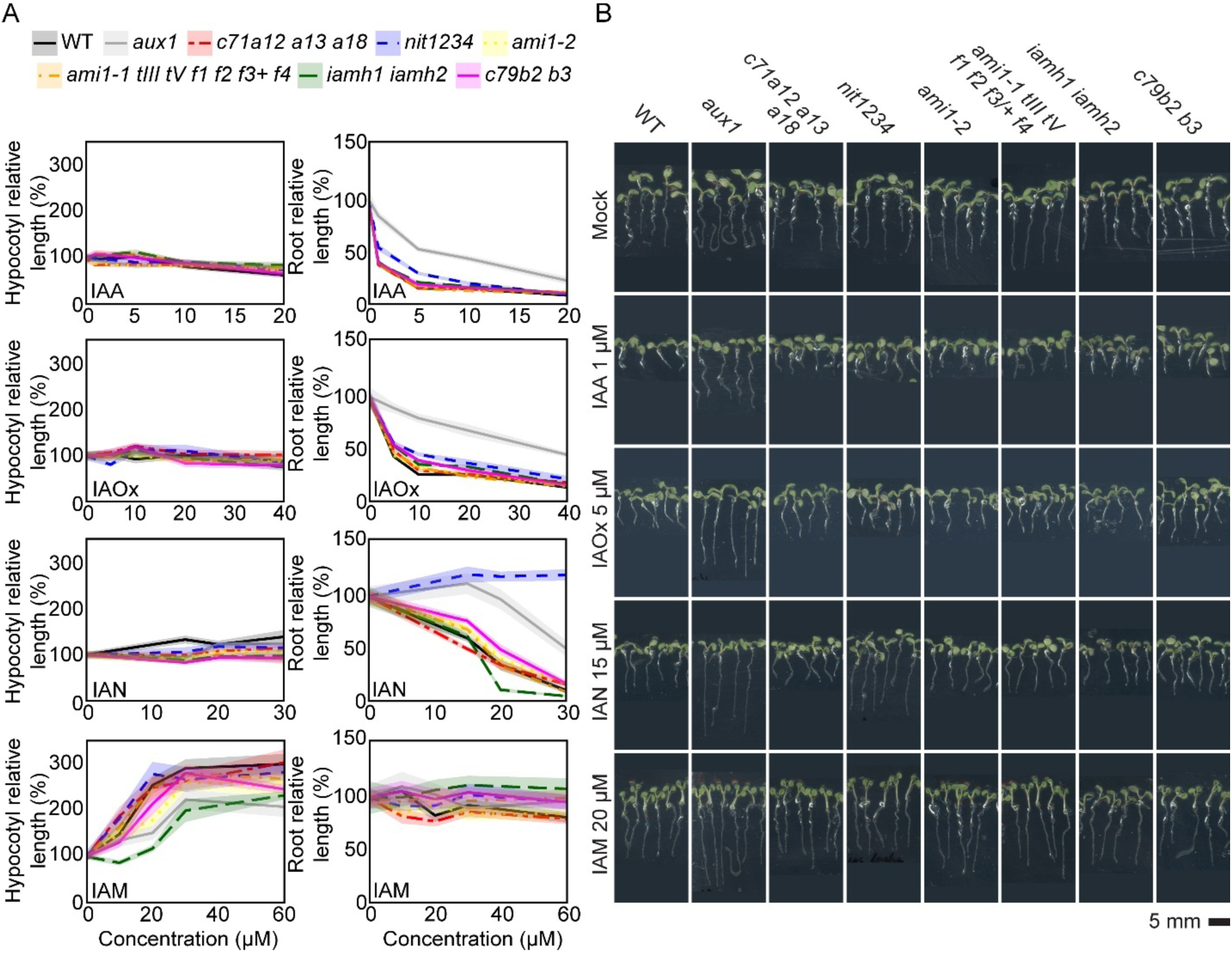
Phenotypes of light-grown mutants impaired in the putative IAOx route challenge the established model of the IAOx pathway. (A) WT and mutant lines were germinated on horizontal plates under continuous light for five days in control media or in media supplemented with the indicated concentrations of IAA, IAOx, IAN and IAM (Supp. Table 2). For each treatment, the “0” concentration contains the equivalent concentration of DMSO as the highest concentration tested for a specific auxin precursor. Root and shoot lengths were measured in ImageJ. Relative organ size at a given concentration for a specific genotype was calculated by dividing the organ size by that in the corresponding control ([metabolite]=0). Average relative organ sizes (lines) and confidence intervals (CI=95%, shades) were plotted using R studio. (B) Photographs of representative plants for one of the concentrations for each compound. These precursor concentrations were chosen as they produce similar organ sizes in the WT as 1µM IAA. WT: wild-type (Col-0), *aux1: aux1-7, c71a12 a13 a18: cyp71a12 cyp71a13 cyp71a18, nit1234: nit1 nit2 nit3 nit4, ami1-1 tIII tV f1 f2 f3/+ f4: ami1-1 toc64-III toc64-V faah1 faah2 faah3/+ faah4, iamh1 iamh2: iamh1-1 iamh2-2, c79b2 b3: cyp79b2 cyp79b3*.

To determine if these phenotypic changes were indeed due to an increase in IAA activity in the treated plants, we examined the effects of these putative auxin intermediates in *aux1-7*, a mutant impaired in the cellular import of IAA and, therefore, compromised in the cell-to-cell movement of this hormone (Pickett et al., 1990; Bennett et al., 1996; Marchant et al., 1999). We found that *aux1* showed partial insensitivity to all precursors (Figure 3; Supplementary Fig. S7), suggesting that the phenotypes observed in WT plants treated with these compounds were, at least to some extent, due to the conversion of these precursors into IAA.

To further verify that the phenotypes observed in WT plants upon exogenous application of these putative IAA precursors were indeed associated with an increase in auxin responses, we examined the activity of the auxin reporter *DR5:GFP* (Figure 4). Three-day-old, etiolated WT seedlings harboring this transcriptional auxin response sensor showed higher *DR5* reporter activity in root tips and root-hypocotyl junctions of plants exposed to IAOx or IAN. Elevated *DR5* activity was also observed in the apical hooks in the presence of all three compounds, with the IAN- and IAM-treated plants also showing partially open hooks concomitant with an especially high activity of *DR5* in these tissues (Figure 4A). When seeds were germinated and grown under continuous light for 5 days, IAOx-treated plants recapitulated the patterns observed in IAA-treated plants, i.e., shorter roots with strong *DR5* activity in the proximal region of the root and root-hypocotyl junction, with a milder induction in the aerial parts (Figure 4B). Although we observed a prominent activation of *DR5* in the aerial parts of both IAN- and IAM-treated seedlings, each compound had different phenotypic signatures. Thus, IAN-treated plants showed shorter roots but no effects on hypocotyl elongation or leaf epinasty, whereas IAM treatment had no effect on roots but dramatically promoted hypocotyl elongation and leaf epinasty (Figure 4B).

**Figure 4.**
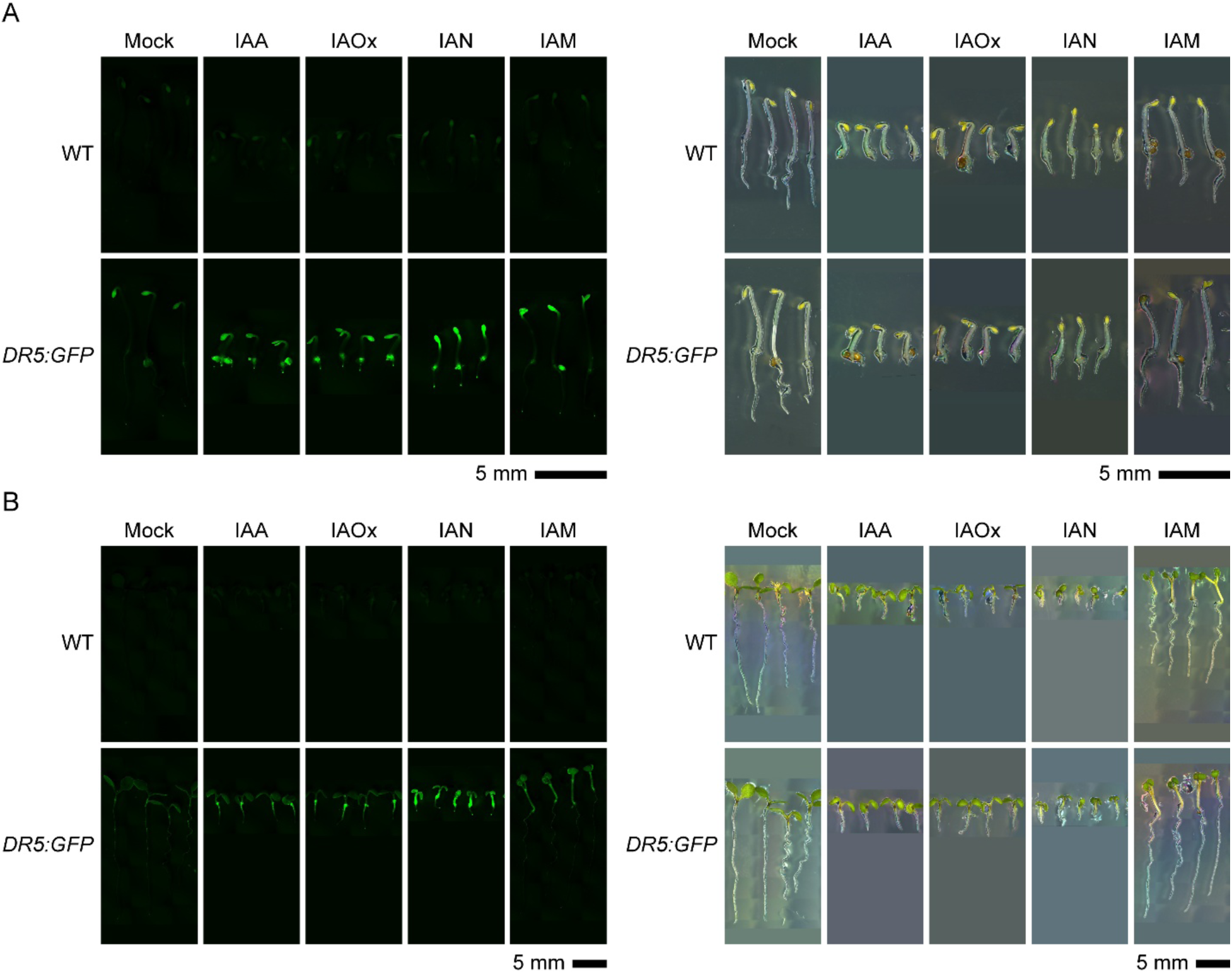
Putative IAOx route intermediates, IAOx, IAN and IAM, induce *DR5:GFP* auxin reporter activity. (A,B) Three-day-old dark-grown seedlings (A) and five-day-old light-grown seedlings (B) were germinated on horizontal AT plates supplemented with the indicated auxin precursors at concentrations empirically determined to be the lowest for seedlings to reach maximum change in organ size (Figure 3A). Dark experiment: 0.3µM IAA, 5µM IAOx, 20µM IAN, 50µM IAM, 0.25µM NAA. Light experiment: 3µM IAA, 20µM IAOx, 30µM IAN, 50µM IAM, 0.5µM NAA.

To rule out the possibility that an apparent increase in *DR5* activity was an artifact due to the smaller size of the organs observed in some treatments, we examined the effects of these compounds on *DR5* activity after a shorter (16-hour) exposure when the phenotypic effects of these treatments on organ size were less prominent (Supplementary Fig. S8). Consistent with what was observed with the longer treatments, 16 hours post transfer of three-day-old, etiolated seedlings to plates containing IAOx or IAN resulted in an increase in *DR5* activity in the roots, while the effects of the IAM treatment was again restricted to the hypocotyls and cotyledons of dark-grown seedlings. Similarly, 16 hours after the transfer of 5-day-old light-grown seedlings to precursor-supplemented plates, we observed an increase in *DR5* activity in the roots of IAOx- and IAN-treated plants and in the hypocotyls and cotyledons of IAM-treated seedlings. These results support the idea that the auxin-related phenotypes triggered by these compounds are indeed due to their conversion into active IAA. Interestingly, each one of these compounds had certain tissue specificity, with IAOx having an effect predominantly in underground tissue (roots), IAM predominantly affecting aboveground tissues (hypocotyls and cotyledons), and IAN acting in both hypocotyls and roots (Figure 4B, Supplementary Fig. S8).

To determine whether or not the conversion of these compounds into IAA requires the activity of the three enzyme families studied here, we examined the phenotypic effects of these compounds in high-order mutants for each of the three gene families in both five-day-old light-grown (Figure 3) and three-day-old dark-grown seedlings (Supplementary Fig. S5). Phenotypic analysis of high-order mutants of the *CYP71A* and *AMI1/TOC64/FAAH* families showed that the response of these mutants to IAOx, IAN, and IAM was not significantly different than that of WT under all of the growth conditions examined (Figure 3; Supplementary Fig. S7). The inability of these high-order mutants to suppress the auxin-like effects of treatments with the proposed IAOx pathway intermediates argues against the previously proposed involvement of these gene families in the conversion of these compounds into IAA *in planta*. In contrast, and as expected from previous work using the single *nit1-3* mutant (Normanly et al., 1997), the roots of the *nit1234* quadruple mutant showed clear insensitivity to the IAN treatment (Figure 3; Supplementary Fig. S7). These results, together with the fact that *nit1234* does not show a consistently altered response to IAA (Supplementary Fig. S7; Supplementary Fig. S10), strongly support the idea that the nitrilase activity is required for the conversion of IAN taken up by the plant into IAA. Furthermore, the profound root insensitivity to IAN observed in *aux1* (Figure 3; Supplementary Fig. S7) is most likely due to the impaired cell-to-cell IAA transport rather than to a putative disruption of the IAN uptake capacities of this mutant, although we have not formally ruled out the latter possibility. Finally, the fact that the *nit1234* responds normally to IAOx suggests that IAN is an unlikely intermediate in the direct conversion of uptaken IAOx into IAA.

In addition to the amidase family described above, two IAM hydrolases, IAMH1 and IAMH2, have been recently implicated in the conversion of IAM into IAA *in vivo* (Gao et al., 2020). Consistent with prior reports, the increase in hypocotyl elongation of five-day-old light-grown seedlings in response to IAM observed in WT plants was significantly attenuated in the double *iamh1 iamh2* mutant (Figure 3). Interestingly, this double mutant responded normally to IAA and the other IAA precursors tested (Figure 3; Supplementary Fig. S7). Following the same logic as used above to interpret the results of the IAN-treated *nit1234* mutant, we conclude that IAM is not a likely metabolic intermediate in the conversion of IAOx and IAN into IAA, as otherwise *iamh1 iamh2* should have been insensitive not only to the IAM but also to the IAOx and IAN supplementation. The lack of IAM insensitivity in *AMI* family mutants, unlike that of *iamh1 iamh2,* indicates that *AMI1/TOC64/FAAH* do not play a prominent role in transforming IAM into IAA in Arabidopsis, and this finding is consistent with the lack of suppression of the high-auxin phenotypes of *sur2* by *ami1-2 toc64-III toc64-V faah1 faah2*. To test if the IAM insensitivity of *iamh1 iamh2* can suppress *sur2*, we introgressed *iamh1 iamh2* into the *sur2* background (Supplementary Fig. S9). Critically, the *iamh1 iamh2 sur2* triple mutant was phenotypically indistinguishable from *sur2*, demonstrating that IAM-to-IAA conversion is not necessary for auxin overproduction in *sur2*.

Unlike the genetically supported NIT- and IAMH-mediated conversion of exogenous IAN and IAM, respectively, into IAA in plants, the lack of genetic evidence for the enzymatic conversion of IAOx into IAA made us wonder whether the auxin-like phenotypes observed in seedlings exposed to exogenous IAOx were caused by spontaneous conversion into IAA—as observed for the IAA precursor IPyA (Stepanova et al., 2008)—or by IAA impurities present in our IAOx stock. To test these possibilities, we first analyzed our lab’s IAOx stocks by high-performance liquid chromatography (HPLC) coupled with electrospray ionization (ESI) and mass spectrometry (MS) (HPLC-ESI-MS). While both *trans-* and *cis*-isomers are present in our IAOx standard, no IAA was detected (Supplementary Fig. S11A). Next, we incubated an IAOx solution (10 ng/uL) in a growth chamber in the light or in the dark for five days, mimicking the experimental conditions used in our phenotyping studies. The LC-MS analysis of these samples showed no conversion of IAOx to IAA regardless of exposure to light (Supplementary Fig. S11B). Therefore, we conclude that the phenotypes observed upon exogenous supplementation of plant growth media with IAOx could not be the consequence of the spontaneous formation of IAA from IAOx *in vitro*.

In summary, our results presented in this section are consistent with the previously proposed idea that IAOx, IAN, and IAM can function as IAA precursors when exogenously provided to plants in the media. They also support the prevalent view that nitrilases and IAM hydrolases play a role in the *in vivo* conversion of exogenously applied IAN and IAM into IAA, respectively. On the other hand, our work questions the existence of the proposed IAOx→IAN→IAM→IAA linear metabolic pathway and the involvement of IAN and IAM as intermediates in the excess auxin production observed in the *sur2* mutant. Our findings instead suggest that each of the three postulated auxin precursors—IAOx, IAN, and IAM—follows an independent route to become an active auxin. Importantly, although NITs and IAMHs are promising enzymes for converting IAN and IAM, respectively, into IAA, no suitable enzyme candidate exists for the transformation of IAOx into IAA *in vivo*. Future genetic and metabolic analysis of *sur2* would be essential for identifying genes coding for such enzymes.

### Mutants in the postulated IAOx pathway do not display prominent auxin defects in classical auxin-related phenotypic assays

The results described so far indicate that the functions of the three gene families examined in this work (*CYP71A, NIT,* and *AMI*) and the *IAMH* family are not required for the excess production of auxin in the *sur2* mutant, nor for normal plant development under standard laboratory growth conditions. This, however, does not rule out the possibility that these genes may participate in the production of auxin under specific growth conditions where a burst of auxin production is required. In fact, *TAA1*, a central component of the IPyA auxin biosynthetic pathway, was originally identified not because of any prominent general developmental defect of the corresponding mutant but due to its altered ethylene (Stepanova et al., 2008) and shade avoidance (Tao et al., 2008) responses, phenotypes that rely on a local increase in auxin production. Thus, we reasoned that to uncover any potential role of the *CYP71A, NIT*, and *AMI* gene families in auxin biosynthesis, the performance of the corresponding high-order mutants should be examined in a battery of phenotypic assays highly sensitive to alterations in the auxin biosynthesis, transport, or signaling pathways. Thus, we first examined the root and hypocotyl sensitivity of the high-order IAOx mutants to the ethylene precursor 1-aminocyclopropane-1-carboxilic acid (ACC) (Merchante and Stepanova, 2017) as it is well known that even mild alterations in the production, transport, or sensitivity to auxin result in a significant reduction in the ACC-triggered growth inhibition. As previously reported (Stepanova et al., 2008), the ACC response of *wei8*, a weak auxin biosynthetic mutant defective in *TAA1*, was normal in the hypocotyls of three-day-old, etiolated seedlings but significantly reduced in the roots (Figure 5A). Furthermore, *sur2* showed WT-level of root sensitivity to ACC, whereas *wei8 sur2* presented an intermediate phenotype, suggesting that production of *sur2-*mediated excess auxin can partially compensate for the *wei8*’s deficit in this hormone. Interestingly, none of the high-order IAOx mutants showed any major defects in ACC sensitivity, and the mild phenotypes initially observed in the *cyp71a12/13/18 sur2* hypocotyls and *nit1/2/3/4 sur2* roots were not reproducible across multiple experimental repetitions (Supplementary Fig. S12) and are thus likely not biologically relevant.

**Figure 5.**
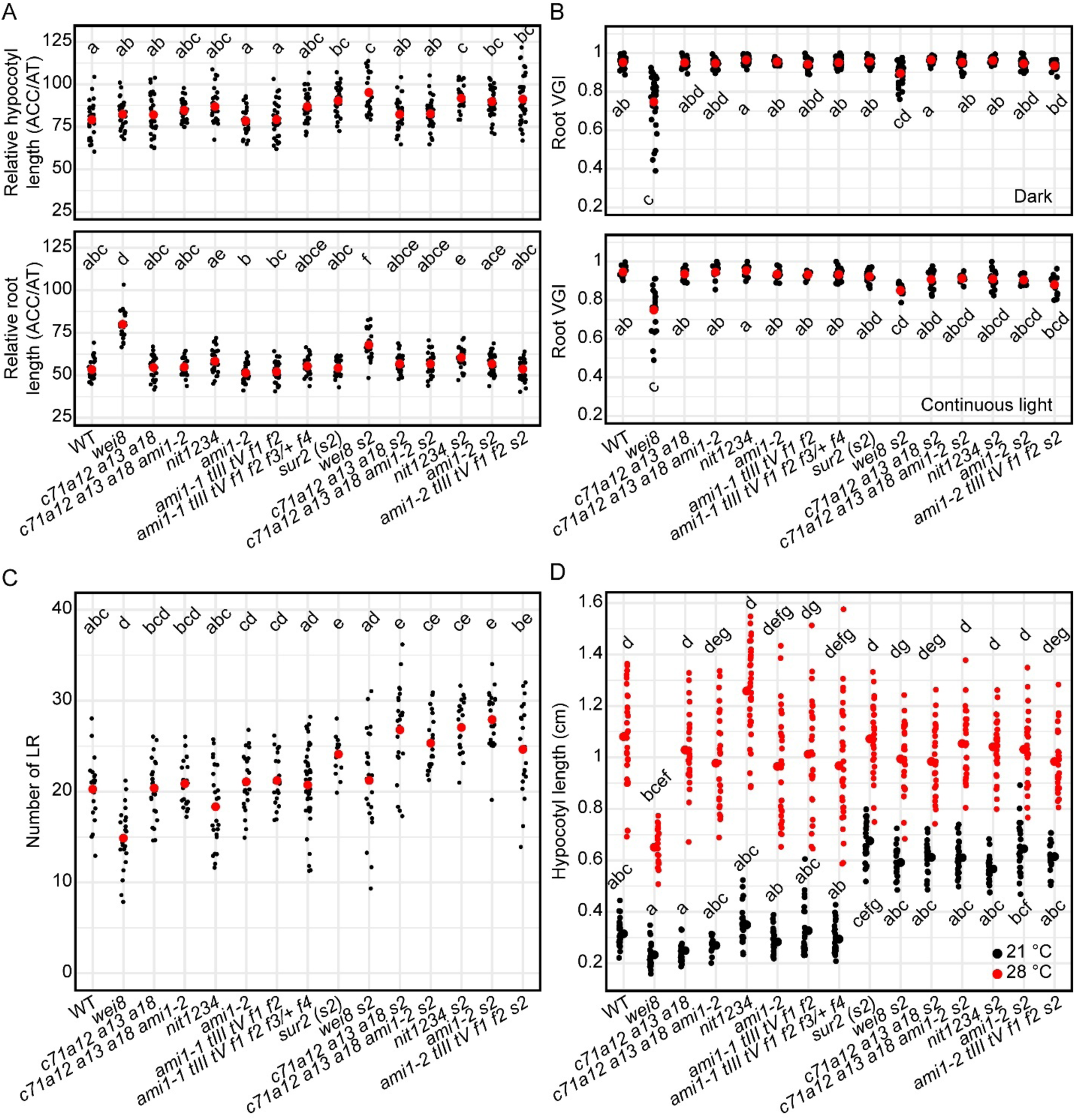
IAOx mutants do not show prominent phenotypes associated with auxin deficiency. (A) Relative growth of hypocotyls (top) and roots (bottom) of three-day-old, etiolated seedlings germinated on horizontal plates in the presence of the ethylene precursor ACC (0.2µM). Relative length was calculated by dividing organ length in ACC by that in the control (AT). (B) Root Vertical Growth Index (VGI, Grabov et al., 2005) of seedlings grown on vertical plates for three days in the dark (top) or for five days in the light (bottom). (C) Number of lateral roots in 10-day-old seedlings grown on vertical plates in continuous light. (D) Heat-induced hypocotyl elongation in 10-day-old, light-grown seedlings germinated on vertical plates. Different letters denote statistically significant differences for α=0.05. WT: wild-type (Col-0), *c71a12 a13 a18: cyp71a12 cyp71a13 cyp71a18, c71a12 a13 a18 ami1-2: cyp71a12 cyp71a13 cyp71a18 ami1-2, nit1234: nit1 nit2 nit3 nit4, ami1-1 tIII tV f1 f2: ami1-1 toc64-III toc64-V faah1 faah2, ami1-1 tIII tV f1 f2 f3/+ f4: ami1-1 toc64-III toc64-V faah1 faah2 faah3/+ faah4, s2: sur2*.

In addition to the altered response to ACC, another hallmark of auxin deficiency is the alteration of gravitropic responses, as previously reported for auxin biosynthesis (*wei8 tar2*; (Stepanova et al., 2008), transport (*aux1*, Pickett et al., 1990; Bennett et al., 1996; Marchant et al., 1999), and signaling (*tir1*, (Hobbie and Estelle, 1994; Ruegger et al., 1998) mutants. Thus, to further explore the possible roles of the gene families postulated to be involved in IAOx-dependent auxin biosynthesis, we examined the root Vertical Growth Index (VGI) as a sensitive measure of gravity responses in both dark- and light-grown seedlings (Marchant et al., 1999; Rahman et al., 2001; Grabov et al., 2005; Stepanova et al., 2008; Rahman et al., 2010; Marquès-Bueno et al., 2021). As expected, we observed an impaired VGI response in the *wei8* mutant, a phenotype that was again (as in the case of ACC response) partially rescued by the excess auxin produced in the *sur2* background. Importantly, none of the high-order IAOx mutants showed significant deviation from the response observed in WT roots (Figure 5B).

Another auxin-regulated process that is highly sensitive to alterations in IAA activity is the formation of lateral roots (Lavenus et al., 2013), with IAA controlling the initiation and emergence processes, as well as patterning (Marchant et al., 2002; Bao et al., 2014; Robbins and Dinneny, 2018). Therefore, we next examined lateral root number in the high-order IAOx mutants. Quantification of the number of lateral roots in 10-day-old seedlings grown vertically under continuous light showed that *wei8* consistently produced fewer and *sur2* more lateral roots than WT, while the rest of the mutants had no appreciable alteration of this phenotype (Figure 5C).

Finally, we examined the rapid hypocotyl elongation of seedlings grown in the light and exposed to high temperatures, as this is also a well-documented, auxin-dependent process (Ruegger et al., 1998). Hypocotyls of WT plants grown at 28°C were approximately four times as long as those of seedlings grown at 21°C (Figure 5D). As expected, a defect in auxin biosynthesis like that of *wei8* translated into a significant reduction in the high-temperature-triggered hypocotyl elongation, a defect that was rescued by the increased auxin production in the *wei8 sur2* plants. In contrast, neither of the high-order IAOx mutants significantly altered the hypocotyl elongation response to high temperature in either the WT or the *sur2* mutant backgrounds.

In summary, none of the highly sensitive auxin-related phenotypic assays we carried out were able to detect significant phenotypic defects of the high-order *CYP71A, NIT,* and *AMI* family mutants. These results thus question the role of these gene families not only in the production of auxin under standard growth conditions, but also in response to abrupt changes in environmental or developmental factors (such as light regimen, temperature fluctuations, gravity vector changes in response to obstacle avoidance, or ethylene buildup in response to stress or soil compactness), as demonstrated here and elsewhere by utilizing classical phenotypic assays known to be highly sensitive to the levels of auxin (Stepanova et al., 2005; Stepanova et al., 2007) that allow to visualize the mild auxin defects of the biosynthetic mutant *wei8* (Stepanova et al., 2008).

### IAOx mutants do not show significant alterations in their IAOx/IAA metabolic profiles

In light of the lack of obvious morphological defects associated with auxin deficiency in the IAOx mutants after employing multiple highly sensitive assays, we decided to further explore the potential role of *CYP71A, NIT,* and *AMI* family genes in auxin biosynthesis by directly quantifying IAA, the IAA precursors anthranilate (ANT), TRP, IPyA, IAOx, IAN, IAM, and the inactivation products indole-3-acetyl-aspartate (IAA-Asp), indole-3-acetyl-glutamate (IAA-Glu), and 2-oxindole-3-acetic acid (oxIAA) (Figure 1). To improve our ability to detect metabolic changes associated with altered IAA biosynthesis, we tested different growth conditions with the goal of identifying those conditions that would result in the strongest phenotypic differences between the WT, the weak auxin-deficient mutant *wei8*, and the *sur2* auxin overproducer. We found that when plants are grown in vertical plates for three days in the dark and then transferred to constant light for four additional days, *sur2* phenotypes such as small epinastic cotyledons and abundant adventitious roots were very prominent (Figure 6). Interestingly, not only these *sur2* defects but also the *wei8* root gravity response and root meristem maintenance were strongly impaired under these growth conditions (Figure 6) relative to plants grown in the same light regimen but in horizontal plates (Figure 2; Supplementary Fig. S5). As expected, the excess auxin production in *sur2* partially rescued the root meristem size and root gravity defects of *wei8*. Even under these growth conditions, where the mild auxin biosynthetic defects of *wei8* were dramatically enhanced, the phenotypes of all of the high-order IAOx mutants were indistinguishable from those of the WT control plants. Nevertheless, we reasoned that these growth conditions were highly sensitive to alterations in the levels of auxin production and, therefore, ideal for examining the ability of the IAOx high-order mutants to alter the IAA metabolic profile in the WT background or to block the expected alterations caused by the *sur2* mutation.

**Figure 6.**
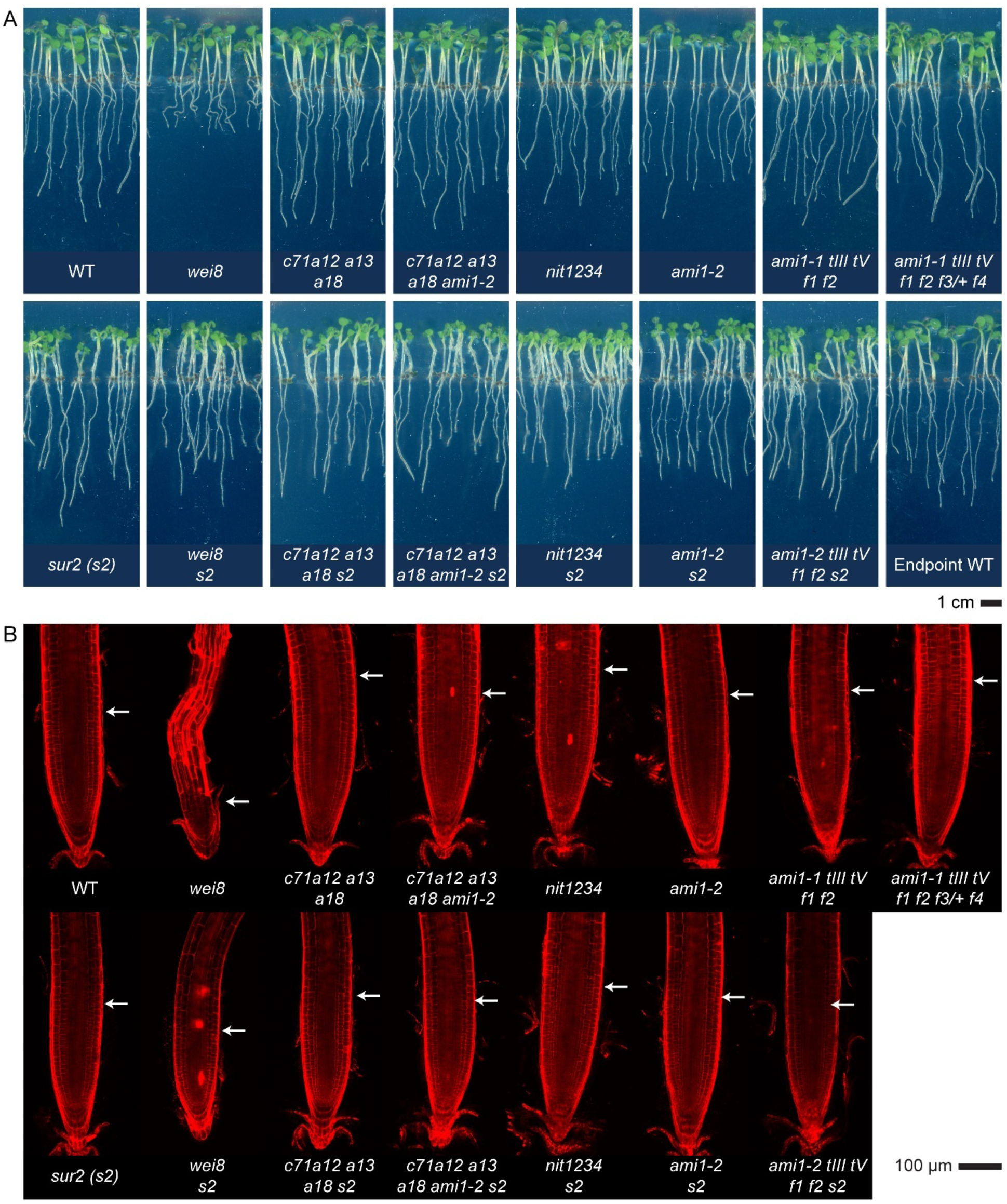
IAOx mutants grown on vertical plates do not show prominent phenotypes. (A) The root growth defect of auxin-deficient *wei8* mutant is prominent when seedlings are grown on vertical plates for three days in the dark followed by four additional days in the light, whereas neither of the IAOx mutants displays prominent defects in these growth conditions. B) Propidium iodide-stained root tips of IAOx mutants are morphologically similar to that of WT plants, unlike that of *wei8* that show meristem degeneration. White arrows point at the end of the root cap, a proxy for the end of the root meristem. WT: wild-type (Col-0), *c71a12 a13 a18: cyp71a12 cyp71a13 cyp71a18, c71a12 a13 a18 ami1-2: cyp71a12 cyp71a13 cyp71a18 ami1-2, nit1234: nit1 nit2 nit3 nit4, ami1-1 tIII tV f1 f2: ami1-1 toc64-III toc64-V faah1 faah2, ami1-1 tIII tV f1 f2 f3/+ f4: ami1-1 toc64-III toc64-V faah1 faah2 faah3/+ faah4, s2: sur2*.

We next employed liquid chromatography coupled to tandem mass spectrometry (LC-MS/MS) to quantify the concentrations of ANT, TRP, IPyA, IAOx, IAN, IAM, IAA, IAA-Asp, IAA-Glu, and oxIAA in WT, selected IAOx high-order mutants, and *wei8* in both WT and *sur2* mutant backgrounds (Figure 7). Due to the large difference in metabolite concentrations (for instance, IAA is only ∼1% of total tryptophan; Supplementary Fig. S13A), to better visualize the data, we normalized each compound to its average WT concentration and represented the fold change by the relative size of the corresponding bubble (Figure 7). Consistent with previous reports (Novák et al., 2012), the concentration of IAOx dramatically increased in the *sur2* mutant. Not surprisingly, none of the high-order IAOx mutants had any effect on the high levels of IAOx accumulating in the *sur2* mutant (Figure 7). Interestingly, the increase of between 60-120-fold in the levels of IAOx observed in all the lines in the *sur2* mutant background did not lead to a higher concentration of IAN or IAM (Figure 7). Moreover, *sur2-*containing lines accumulated lower amounts of IAM than equivalent mutants without *sur2*, with the lowest IAM levels seen in the *aJ tIII tV f1 f2 s2* mutant, which, based on the currently accepted models, would be expected to accumulate the highest concentration of this precursor by blocking the conversion of IAM into IAA.

**Figure 7.**
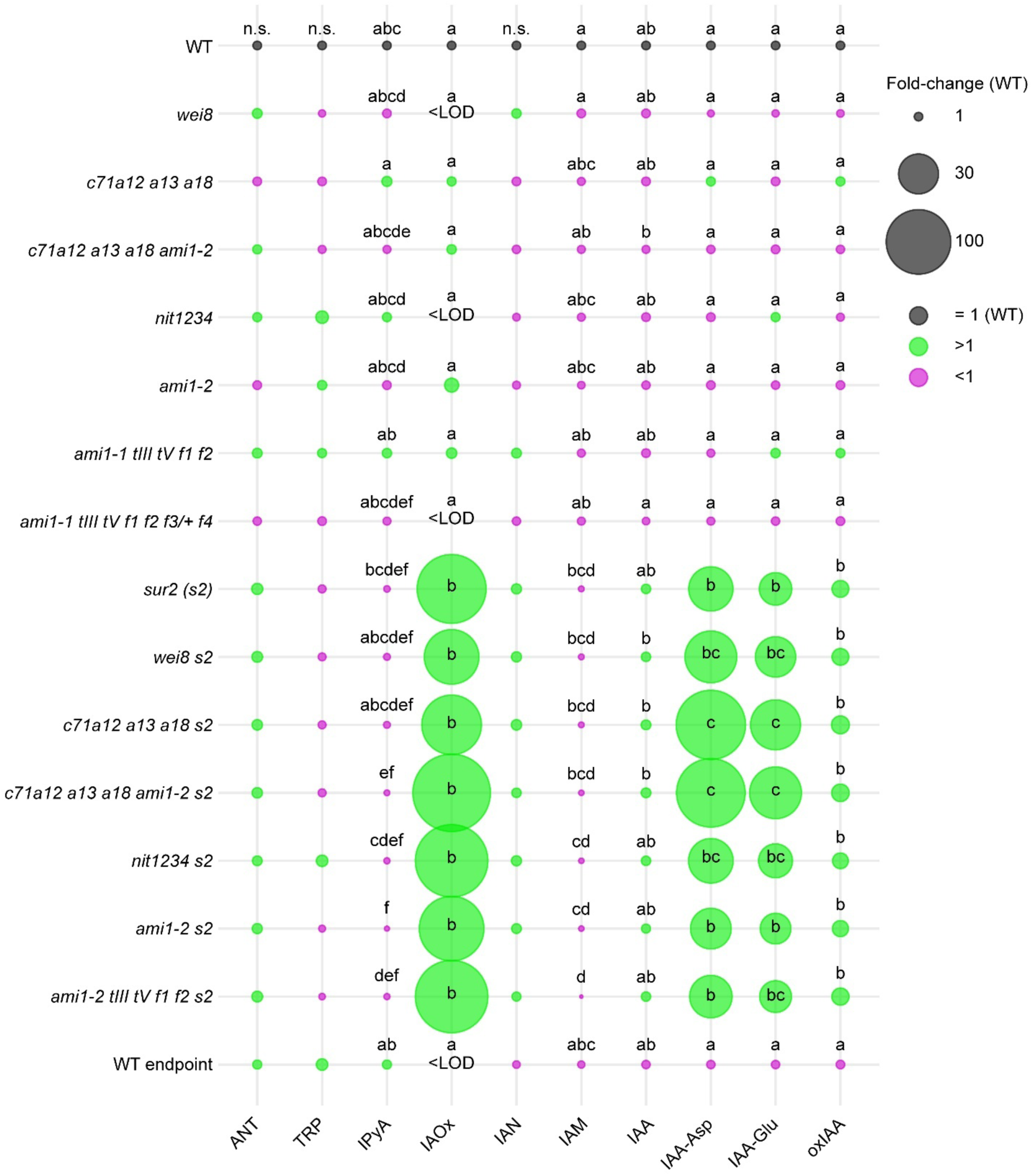
Metabolic quantification in IAOx mutants rules out a prominent role of *CYP71*, *NIT*, and *AMI1/TOC/FAAH* gene families in auxin biosynthesis. Concentrations are normalized to WT values for each metabolite. Bubble sizes are proportional to the concentration fold changes for each mutant compared to WT. Fold change of >1 is shown in green, and that of <1 is in magenta. Different letters denote statistically significant differences between mean values of metabolite concentrations (log10 (pmol g FW^-1^); Supplementary Fig. 9A). WT: wild-type (Col-0), *c71a12 a13 a18: cyp71a12 cyp71a13 cyp71a18, c71a12 a13 a18 ami1-2: cyp71a12 cyp71a13 cyp71a18 ami1-2, nit1234: nit1 nit2 nit3 nit4, ami1-1 tIII tV f1 f2: ami1-1 toc64-III toc64-V faah1 faah2, ami1-1 tIII tV f1 f2 f3/+ f4: ami1-1 toc64-III toc64-V faah1 faah2 faah3/+ faah4, s2: sur2*.

In addition to higher IAOx concentrations in *sur2-*containing mutant lines, we observed a mild increase of auxin concentration, accompanied by a strong increase in the levels of IAA conjugation and oxidation products, metabolic changes also reported previously for *sur2* (Novák et al., 2012; Pěnčík et al., 2013). These *sur2* metabolic changes were not affected by mutating any of the three IAOx gene families studied here. Finally, Principal Component Analysis (PCA) of the IAA-related metabolic profiles showed that the main factor affecting the metabolic profiles in all our experiments is the *sur2* mutation, while none of the mutations in the IAOx gene families that were supposed to work downstream of *SUR2* in the production of IAA had any effect on the *sur2* metabolic profile (Supplementary Fig. S13B).

Together, these results are consistent with our genetic and pharmacological observations (Figure 3; Supplementary Fig. S7) and strongly suggest that the previously proposed metabolic route for the production of IAA from IAOx via IAN and IAM, as well as the corresponding genetic pathway, need to be thoroughly reevaluated.

## Discussion

Several routes for the production of the key auxin IAA have been proposed (Figure 1) (Zhao, 2010). However, only the IPyA pathway has gathered enough experimental support to be generally accepted as the predominant source of IAA in all plants investigated (Zhao et al., 2001; Stepanova et al., 2008; Tao et al., 2008; Yamada et al., 2009; Stepanova et al., 2011; Zhou et al., 2011; Eklund et al., 2015). Two key factors hindered our ability to conclusively test the relevance of alternative IAA production routes. First, there was a lack of mutants that block the activity of entire multigene families coding for the enzymes thought to be involved in these metabolic pathways. Second, it is difficult to rule out possible biosynthetic pathways based on the absence of phenotypic defects in the corresponding knockout mutants, especially since these mutants can, inevitably, be examined only under a limited set of experimental conditions. To address these considerations, we took three complementary approaches. We began by generating high-order mutants knocking out all putative IAOx pathway genes in the three selected multigenic families that have been previously proposed to catalyze the conversion of IAN, IAM, or IAOx into IAA. We then examined the phenotypes of these whole-gene family knockouts under normal growth conditions as well as after specific treatments known to trigger phenotypic changes that are highly sensitive to auxin level disturbances. We did this work not only in the WT but also in the *sur2* mutant background, where the IAOx pathways were believed to be hyperactive. Finally, in addition to this extensive phenotypic characterization, we also evaluated the metabolic profiles of the generated high-order mutants for a battery of IAA precursors and degradation products.

Our results conclusively show that the functions of the three gene families of *CYP71A*s, *NIT*s, and *AMI*s tested in this study are not required for the normal development of Arabidopsis plants under standard laboratory conditions. Importantly, the inability of these high-order mutants to block the high auxin phenotypes of *sur2* or to alter the characteristic metabolic profile of IAA-related compounds in the *sur2* mutant completely rules out any prominent role for these genes in the route of auxin production activated by the *sur2* mutation. These results not only disprove the previously proposed genetic pathway for the production of excess auxin in *sur2* (Figure 1) but also cast doubts about the postulated metabolic pathway where the high levels of IAOx in *sur2* were thought to be converted into IAA via IAN and IAM. However, although we can discount the involvement of *CYP71A*, *NIT*, and *AMI* genes in the IAOx pathway and *sur2*-mediated auxin biosynthesis, we cannot rule out the involvement of IAOx→IAN→IAM→IAA route itself in IAA production of excess auxin in *sur2.* It is still theoretically possible that these metabolic reactions are catalyzed by other, yet unknown enzymes. However, the possibility that this sequence of reactions is responsible for the high IAA levels operating in *sur2* is further weakened by our results from the treatment of the high-order mutants with the different auxin precursors, as summarized in Figure 8. We showed that WT plants treated with each of the IAA precursors, IAOx, IAN, and IAM, displayed, as expected (Normanly et al., 1997; Buezo et al., 2019; Gao et al., 2020; Roman et al., 2023), auxin-related phenotypes (Figure 4, Supplementary Fig. S8). Elimination of the *NIT* gene family function rendered clear resistance to IAN, indicating that nitrilase activities encoded by this gene family are required for the production of IAA from IAN taken up by the plant. Importantly, the *nit1/2/3/4* mutant showed normal response not only to IAA but also to IAOx. This was an unexpected result as, in the previously proposed metabolic pathway, IAN is an intermediate in the production of IAA from IAOx, and therefore, blocking the conversion of IAN into IAA should have also prevented the formation of IAA from IAOx via IAN. Furthermore, in disagreement with a mild insensitivity to IAM reported for the same *ami1-1* and *ami1-2* single mutants (Pérez-Alonso et al., 2020), our high-order *AMI* family mutant responded normally to all IAA precursors tested, including IAM, questioning the prominent role of AMI in the production of IAA *in vivo*. Conversely, the recently published *iamh1 iamh2* mutant showed clear resistance to IAM, as previously reported (Gao et al., 2020), firmly implicating the two *IAMH1 IAMH2* genes in the conversion of exogenous IAM to IAA in seedlings. The response to IAN and IAOx of the *iamh1 iamh2* mutant was, however, undisguisable from that of the WT plants. These results, again, argue against the previously proposed metabolic pathway model where both IAOx and IAN function as IAA precursors upstream of IAM. This is because if the model was accurate, blocking the conversion of IAM into IAA in the *iamh1 iamh2* mutant should result in resistance to both IAOx and IAN, which is not the case. Furthermore, our results indicate that neither of the three gene families investigated here, *CYP71A*s, *NIT*s, and *AMI*s, is involved in the excess auxin production in *sur2*.

**Figure 8.**
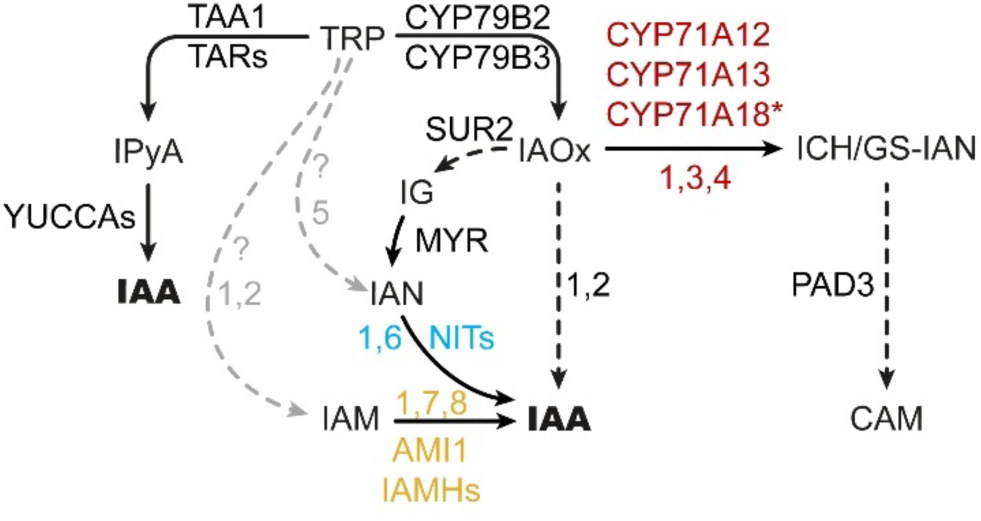
Revised model of TRP-dependent auxin biosynthesis. Black and gray arrows represent high- and low-confidence enzymatic conversions, respectively. Solid arrows indicate single-step catalytic reactions, whereas dashed arrows represent multi-step conversions. Numbers next to arrows correspond to references supporting the connections: 1 - This work; 2 - Sugawara et al., 2009; 3 - Muller et al., 2015; 4 - Mucha et al., 2019; 5 - Zhao et al., 2002; 6 - Normanly et al., 1997; 7 – Pérez-Alonso et al., 2020; 8 - Gao et al., 2020. Trp: tryptophan, IPyA: indole-3-pyruvic acid, IAA: indole-3-acetic acid, IAOx: indole-3-acetaldoxime, IAN: indole-3-acetonitrile, ICH: indole-3-cyanohydrin, GS-IAN: glutathione-indole-3-acetonitrile, IAM: indole-3-acetamide, IG: indole glucosinolate, CAM: camalexin; TAA1: TRP AMINOTRANSFERASE OF ARABIDOPSIS1 (WEI8), TAR: TAA1-RELATED (TAR1, TAR2), NITs: NITRILASE (NIT1-NIT4), IAMH: IAM HYDROLASE (IAMH1, IAMH2), MYR: MYROSINASE, PAD3: PHYTOALEXIN DEFICIENT3, ?: unknown enzyme(s). **Note 1**: *cyp71a12 cyp71a13* double mutant is CAM-deficient (Muller et al., 2015) but not insensitive to IAOx; *cyp71a12 cyp71a13* is GS-IAN deprived (Muller et al., 2015) but not IAN-deprived; and according to Mucha et al., 2019, CYP71A12 and CYP71A13 make indole-3-cyanohydrin. Since we did not quantify CAM-related compounds downstream of the CYP71 activity, and due to the absence of phenotype in *cyp71a12 cyp71a13 cyp71a18* triple mutant generated in this work, the role of CYP71A18 in CAM biosynthesis is yet to be determined, hence showing a star (*). **Note 2**: *cyp79b2 cyp79b3* double mutant did not have detectable levels of IAN according to Sugawara et al. (2009), but according to Zhao et al. (2002), *cyp79b2 cyp79b3* did have detectable IAN. We found that the IAOx->IAN conversion is unlikely to be prominent in auxin biosynthesis, so IAN must have another source. The consensus in the field is that IGs are converted into IAN by means of myrosinases. However, (Sugawara et al., 2009; Novák et al., 2012) reported that *sur2* and *sur1* (impaired in IG biosynthesis) have smaller but detectable amounts of IAN compared to WT, suggesting that IAN does not fully depend on IGs. **Note 3**: IAM is detected in the *cyp79b2 cyp79b3* mutant (Sugawara et al., 2009) suggesting that IAM production is not fully IAOx-dependent. If auxin-like effects of exogenous IAOx or IAN depended on their conversion into IAM, *iamh1 iamh2* double mutant would be partially insensitive. Because *iamh1 iamh2* mutant is as sensitive to IAOx and IAN as WT, IAM may originate from another source.

One may argue that a possible reason for the lack of the *sur2* defect suppression by the mutants in the *CYP71A*, *NIT*, and *AMI* gene families characterized herein is that these putative IAOx pathway genes operate in different cell types from those where *SUR2* is expressed. In that scenario, other, yet unknown, enzymes would be responsible for the conversion of IAOx into auxin in *sur2*. However, there is at least a partial overlap in the cell types where *SUR2* and these other gene families are detected (Supplementary Fig. S14), so at least partial suppression of the *sur2* auxin defects would have been expected by the high-order IAOx family mutant combinations. One could also argue that the residual expression of *FAAH1* and *FAAH4,* as well as the remaining WT copy of *FAAH3,* in the high-order mutant *ami1-1 toc64-III toc64-V faah1 faah2 faah3/+ faah4* are the reason for the lack auxin-deficient phenotypes in this mutant and for the inability of the *ami1-2 toc64-III toc64-V faah1 faah2* to suppress *sur2*. In a scenario where the spatiotemporal distribution of FAAH1, FAAH4, and FAAH3 proteins coincides with an overaccumulation of IAM as their low-affinity substrate (Pollmann et al., 2006), it could theoretically be possible that one or more of these FAAH members catalyze the IAM-to-IAA conversion. However, the low sequence identity that FAAH and AMI1 proteins share (Supplementary Fig. 1A), the lack of prominent accumulation of IAM in our high-order mutants (Figure 7), along with the reported very low *in vitro* activity of FAAH1 against IAM as a substrate yet its strong activity against fatty acid amides (Shrestha et al., 2003; Pollmann et al., 2006; Shrestha et al., 2006; Kim et al., 2009; Khan et al., 2017), renders this hypothetical scenario also unlikely.

Overall, our findings suggest that although exogenously provided IAOx, IAN, and IAM can indeed serve as precursors to IAA, the metabolic relationships previously proposed to link these compounds to the production of IAA in *sur2* should be reconsidered. The results from our targeted metabolic analysis are also inconsistent with what would be expected if the previously proposed metabolic and genetic pathways were correct. Thus, we did not observe any alteration in the endogenous levels of the examined metabolites, including IAA, IAA inactivation products, IAOx, IAN, and IAM, in our high-order mutants. It could be argued, however, that this is not strong evidence against the involvement of these gene families in the production of IAA under normal conditions, where the IAOx pathway may not be active. However, it would be more difficult to reconcile the lack of changes in the metabolic profiles of the IAA biosynthetic pathway when comparing *sur2*, where the IAOx pathway is supposed to be hyperactive, and our high-order IAOx mutants in the *sur2* mutant background.

Several key questions arise from the results presented in this study. How is excess IAA produced in the *sur2* mutant? Do the IAA production mechanisms activated in the *sur2* mutant play a physiological role in plants, and if so, are they necessarily restricted to *Brassicaceae* species? Finally, are *NIT*s and *IAMH*s involved in a yet uncharacterized IAA biosynthetic pathway? Even though we are currently unable to provide answers to these questions, this work has identified knowledge gaps that were previously ill-defined. Thus, for example, a new hypothesis about how excess IAA is produced from extra IAOx in *sur2* needs to be formulated and tested. Towards that goal, our lab is currently carrying out genetic suppressor screens to identify mutant genes capable of masking the high-auxin phenotypes of *sur2*. Similarly, we have conducted a chemical screen to identify small molecules capable of suppressing the high *DR5:GUS* activity in the *sur2* mutant. Finally, we are performing a non-targeted metabolic analysis of *sur2* to identify compounds potentially involved in the conversion of IAOx into IAA in plants. Deciphering the mechanism by which excess auxin is produced in *sur2* would open the door to investigating whether these mechanisms are also active in WT plants under specific conditions or in response to certain biotic or abiotic factors. One may argue that since IAOx and the corresponding biosynthetic enzymes CYP79B2 and CYP79B3 have not been found outside the *Brassicaceae* family, any potential physiological relevance of this pathway found in *Arabidopsis* would be restricted to a small group of plants. However, the discovery of phenylacetaldoximes and their corresponding biosynthetic enzymes in both monocot and dicot leaves (Perez et al., 2021), along with the observation that IAOx triggers auxin signaling and *sur2*-like phenotypes in a *Fabaceae* species, *Medicago truncatula* (Buezo et al., 2019; Roman et al., 2023), and can be converted to IAA in other non-*Brassicaceae* species (Rajagopal and Larsen, 1972), opens the possibility that insights gained about the IAOx pathways in Arabidopsis could have implications beyond *Brassicaceae* family.

## Materials and methods

### Generation of mutant lines

T-DNA lines and their respective insertion sites are depicted in Supplementary Fig. S2. Higher-order mutant combinations were obtained by crossing, and the desired mutant combinations were identified by genotyping. All the mutant lines generated in this work and their sources are listed in Supp. Table 1. Primer sequences and primer combinations used for genotyping are available in Supp. Table 2 and 3, respectively. *CYP71A* triple KO mutant (*cyp71a12/a13/a18*) was obtained by crossing the previously described *cyp71a12^TALENs^/a13* double mutant (Müller et al., 2015) to *cyp71a18* available from Arabidopsis Biological Resource Center (ABRC). *nit3* and *nit4* mutants were also ordered from ABRC and intercrossed. Due to the tight linkage between *NIT1*, *NIT2,* and *NIT3*, CRISPR/Cas9 genome editing was employed to obtain higher-order *nit* mutants. The guide RNA (gRNA) (5’-ATTGGAAAACTCGGTGCTGC-3’) was cloned into pDONR207 by site-directed, ligase-Independent mutagenesis (SLIM) (Chiu et al., 2004) using the following primers: SLIM_F GTTTTAGAGCTAGAAATAGCAAG, SLIM_R CAATCACTACTTCGACTCT, NITDFor_tailed GCAGCACCGAGTTTTCCAATGTTTTAGAGCTAGAAATAGCAAG, and NITDRev_tailed ATTGGAAAACTCGGTGCTGCCAATCACTACTTCGACTCT. Briefly, two separate inverse PCR amplifications (SLIM_F+NITDRev_tailed and NITDFor_tailed+SLIM_R) were pooled, treated with *DpnI*, melted, reannealed to obtain a linear vector with 20-mer sticky ends, and directly transformed into *Escherichia coli* (*E. coli*). The pDONR207_gRNA plasmid was extracted using alkaline lysis and the gRNA was moved into the Gateway-compatible binary vector pMTN3164 (Denbow et al., 2017) by LR reaction using manufacturer-recommended protocols (Thermo Fisher Scientific). gRNA integrity was confirmed by Sanger sequencing using primers CAS9zn_GW_For TACAACAGTCTTGACACAGTCTCCC and CAS9zn_GW_Rev AGATAGCCCAGTAGCTGACATTCAT.

Arabidopsis *nit3 nit4* double mutant plants were grown in soil under long-day photoperiod, 16-h light/8-h dark, at 20°C. Transformation was carried out using the floral dip method (Clough and Bent, 1998), using *Agrobacterium tumefaciens C58* harboring the gRNA in pMTN3164. T1 transformants were selected on half-strength Murashige and Skoog medium supplemented with 20 μg/ml hygromycin under the described growth conditions. After two weeks, transformed seedlings with true leaves and roots were transferred to soil. A single rosette leaf per T1 plant was harvested, its genomic DNA extracted, and genotyping PCRs performed to confirm the presence of *Cas9* (using internal primers CAS9 inter For: TCCACTGGCTAGAGGCAACT, and CAS9 inter Rev: GCGATATGCTCGTGAAGTGA) and somatic CRISPR-induced deletions in the *NIT2-NIT1* region (Suppl. Table 2-3). The size of the PCR product of the WT *NIT2-NIT1* region was 4.4 Kb, whereas the size of the PCR product of the deleted region was 0.8 Kb. Similarly, T2 plants were genotyped to identify plants without the *Cas9* construct but that harbored the *NIT2-NIT1* deletion. In T3 plants, the homozygosity of the deleted region was confirmed by PCR. Sequencing of the deleted region was performed to rule out the possibility of the generation of a potential chimeric protein resulting from the fusion of the 5’ end of *NIT2* and the 3’ end of *NIT1* genomic regions. The sequencing revealed an insertion of an additional G (Supplementary Fig. S3) that produced a frameshift leading to a premature stop codon 10 amino acids downstream of the targeted site. The selected *Cas9*-free homozygous *nit1 nit2 nit3 nit4* plants were then crossed to *nit3 nit4 sur2* plants to obtain the final *nit1 nit2 nit3 nit4 sur2* quintuple mutant.

*AMI1* gene mutants, *ami1-1* (SALK_069970) and *ami1-2* (SALK_019823), were obtained from ABRC. Mutants in the *TOC64* and *FAAH* family members were obtained from other laboratories: *ami1-1 toc64-III-1 toc64-V-1* (Aronsson et al., 2007) was a gift from Paul Jarvis (Oxford University) and Henrik Aronsson (University of Gothenburg), and *faah1 faah2* (Keereetaweep et al., 2013) was provided by Kent Chapman (University of Northern Texas) and Elison Blancaflor (Noble Foundation). Mutants *aux1-7* (Pickett et al., 1990), *wei8-1* (Stepanova et al., 2008), *sur2* (Delarue et al., 1998; Stepanova et al., 2005), *cyp79b2 b3* (Zhao et al., 2002), and *cyp79b2 b3 s2* (Stepanova et al., 2011), as well as *YUCCA1* overexpression line (*YUCox*) (Zhao et al., 2001) have been previously described. Mutant *iamh1-1 iamh2-2* (Gao et al., 2020) was a gift from Prof. Hiroyuki Kasahara (Tokyo University of Agriculture and Technology).

### Growth conditions

Seeds were surface-sterilized using a solution consisting of 50% commercial bleach supplemented with 0.01% Triton X-100 for 5 minutes, washed with sterile de-ionized water four times, and stratified for three days at 4°C prior to starting any assay. For ACC sensitivity assay, seeds were resuspended in sterile 0.6% low-melting-point (LMP) agarose, sowed in aseptic conditions on the surface of sterile AT media (4.33 g/L Murashige and Skoog salts, 10 g/L sucrose, pH 6.0 adjusted with 1M KOH, 6 g/L Bacto Agar) and AT supplemented with 0.2 µM ACC. Germination was induced under ambient light for 2 hours at room temperature and the plates were transferred to the dark at 22°C and kept horizontally. After 72 hours, seedlings were individually transferred onto another plate for imaging. For quantification of seedling sensitivity to IAA (Sigma-Aldrich) and IAA precursors IAOx (Ambeed), IAN (Sigma-Aldrich), IAM (Sigma-Aldrich), seed sowing and germination on supplemented plates were identical to those described for ACC, with the assays run in horizontal plates under a range of precursor concentrations, as shown in Supp. Table 4, either under continuous LED light (70-100 µmol m^-2^ s^-1^; 2x 6000K Kihung T8 LED integrated fixture 40W + 1x FULL SPECTRUM Monios-L LED grow light full spectrum 60W) for 5 days or in the dark for 3 days.

For *DR5:GFP* reporter activity assays, seed sowing and germination on supplemented plates were identical to those described for the IAA precursor treatments, selecting the concentrations of IAA intermediates that produce comparable degree of phenotypic responses across all precursor treatments. For plant imaging, seedlings germinated on supplemented plates were transferred to plain 0.8% (w/v) Bacto Agar plates and imaged using Leica Thunder Imager M205FA equipped with Leica DMC6200 color camera (bright field) and Leica DFC9000 sCMOS camera (fluorescence). For shorter 16-hour precursor treatments, seedlings were sowed and germinated on horizontal plates, as described for ACC assays, but on control (AT) plates without ACC supplementation. Dark-germinated seedlings were grown for ∼2.5 days, transferred onto supplemented plates, and grown vertically for additional 16 hours in the dark before imaging to complete 72 hours. Light-germinated seedlings were grown for ∼4.5 days, transferred onto supplemented plates, and grown vertically for additional 16 hours under continuous LED light before imaging.

To assess VGI (Grabov et al., 2005), seed sowing and germination on supplemented plates were identical to those described for ACC. Seeds were resuspended in 0.2% (w/v) LMP agarose, and sowed on square AT plates for vertical growth (10 g/L Bacto Agar) in a line containing 20-30 seeds. Germination was induced as described above and plates were placed vertically at 22°C in the dark or under continuous LED light for 5 days prior to image acquisition. For the quantification of lateral roots, seeds were sowed and germinated as described for VGI and grown vertically under continuous light for 10 days prior to imaging. Counting of emerged lateral roots was assisted by a Nikon SMZ645 stereo microscope, and small bulges without epidermal opening were not counted. For heat-induced hypocotyl elongation tests, seed sowing and germination were identical to those described for the VGI experiment, and assays were performed as described (Zhu et al., 2021) in Percival I-36LL chambers with INTELLUS control system under constant fluorescent light (70 µmol m^-2^ s^-1^). For metabolic profiling, seeds were surface-sterilized, sowed and germinated as described for VGI, and grown in the dark for three days followed by four days under continuous LED light. 7-day-old seedlings were harvested and flash-frozen in liquid nitrogen, ground manually using a liquid nitrogen-prechilled mortar and pestle, and weighed prior to sample lyophilization.

To image adult plants grown in soil, seeds were sowed and germinated as described for the VGI assay. Plates were then placed vertically in the dark for three days at 22°C and transferred to continuous light for four more days prior to imaging. Then, seedlings were transferred to soil (1:1 ratio of SunGro professional growing mix and Jolly Gardener Pro-Line C/B growing mix) to standard 4×6 flats (Greenhouse Megastore), at 5 seedlings per each pot, 4 pots per genotype. Pot positions were semi-randomized, with the pots periodically reshuffled to minimize positional effects. Plants were grown for two more weeks under white light using fluorescent bulbs (EIKO F54T5/HO/850 TCLP 1C2) in long-day conditions (16-h light/8-h dark) at 22°C prior to imaging using an OLYMPUS PEN Lite E-PL6 camera.

### Quantification of auxin, auxin precursors, and auxin metabolites

Auxin metabolite profiles were analyzed using liquid chromatography–tandem mass spectrometry (LC-MS/MS) following the method described by (Novák et al., 2012). Briefly, approximately 10 mg of fresh weight per sample were lyophilized and extracted with 1 mL of cold 50 mM phosphate buffer (pH 7.0) containing 0.1% sodium diethyldithiocarbamate and a mixture of stable isotope-labelled internal standards. First, 500 µL portion of the centrifuged extract was acidified to pH 2.7 with HCl and purified by solid-phase extraction (SPE) using Oasis^TM^ HLB columns (30 mg, 1 mL; Waters, USA). Second, 500 µL portion was derivatized with cysteamine, acidified to pH 2.7 with HCl, and purified by SPE to determine IPyA. Following elution, all samples were evaporated under reduced pressure, reconstituted in 10% aqueous methanol, and analyzed using I-Class UHPLC system (Waters, Milford, CT, USA) equipped with Kinetex C18 column (50 mm×2.1 mm, 1.7 µm; Phenomenex) and coupled to a triple quadrupole mass detector (Xevo TQ-S; Waters, USA).

### Assessment of spontaneous conversion of IAOx into IAA

Chemical standards of IAA and IAOx were analyzed on an Agilent G6530A QTOF LC/MS instrument. A Zorbax Eclipse Plus C18 column (3 x 100 mm, 1.8 μm) was used with a binary gradient of 0.1% (v/v) formic acid in water (solvent A) and 0.1% (v/v) formic acid in acetonitrile (solvent B) at a flow rate of 0.6 mL min^-1^. The gradient started at 5% solvent B for 1 min, followed by a linear increase to 65% B over 6 min, then to 95% B over 1 min, and held at 95% B for 2 min. The acquisition of mass spectra was done in the positive ionization mode with the following parameters: drying gas temperature, 300 °C; drying gas flow rate, 7.0 L min^−1^; nebulizer pressure, 40 psi; sheath gas temperature, 350 °C; sheath gas flow rate, 10.0 L min−1; Vcap, 3500 V; Nozzle Voltage, 500 V; Fragmentor, 150 V; Skimmer, 65.0 V; Octopole RF Peak, 750 V.

### RNA extraction and qPCR

Seeds from each genotype were sown on 8 g/L agar plates and incubated horizontally under continuous light at 22 °C for 10 days prior to tissue collection. Seedlings from two plates per genotype were harvested and flash frozen in liquid nitrogen. Frozen seedlings were ground in a liquid nitrogen-prechilled mortar, and 100 mg of pulverized tissue from each genotype were mixed with 1 mL of Trizol LS reagent (Invitrogen) per sample and phase separated using chloroform. RNA was extracted from the upper aqueous phase using the Qiagen RNeasy Kit following the manufacturer protocol (Qiagen) with an on-column DNase treatment for 15 minutes at 30 °C (Qiagen). First-strand cDNA synthesis was performed on 100 ng of total RNA using an oligo-d(T)18 primer and 2 µL of SuperScriptIII reverse transcriptase (Invitrogen). Reverse transcription reactions were incubated for 1 hour at 50 °C, 15 minutes at 55 °C, followed by 15 minutes at 70 °C for enzyme deactivation. The product of each reaction was diluted 5-fold and 3 µL of the dilution were used as a template for RT-qPCR with a SYBR Green PCR Mastermix (Applied Biosystems) in a StepOnePlus Real-Time PCR system (Applied Biosystems). A default program of 10 min at 95 °C, followed by 40 cycles of {15 sec at 95 °C, 30 sec at 60 °C} was utilized.

To compare levels of expression across different genotypes, primers were designed to amplify a <250bp region at the 3’ end of mature transcripts. For all gene targets, one or both primer sequences overlapped an exon-exon junction to selectively amplify from cDNA. Primer efficiency was determined for each primer pair using a serial dilution of WT (Col-0) cDNA, with three replicates per sample. Given similar primer efficiencies, relative gene expression was evaluated using the ΔΔCт method (Livak and Schmittgen, 2001). For all targets, RT-qPCR was performed using three biological replicates per genotype. Primers used in expression analysis of intronic alleles by qPCR are available in Supp. Table 5, and annealing sites are depicted in Supplementary Fig. S2.

### Image acquisition and statistical analysis

For the analysis of seedling morphometric traits (ACC-mediated hypocotyl and root shortening, VGI, heat-induced hypocotyl elongation, and counting of emerged lateral roots), seedling images were acquired using an Epson Perfection V600 Photo scanner and analyzed with FIJI/ImageJ. Data plotting and statistical analysis were performed using *ad-hoc* R studio pipelines.

### AGI gene ID

AT1G70560 (*WEI8/TAA1*), AT4G31500 (*SUR2*), AT2G30750 (*CYP71A12*), AT2G30770 (*CYP71A13*), AT1G11610 (*CYP71A18*), AT3G44310 (*NIT1*), AT3G44300 (*NIT2*), AT3G44320 (*NIT3*), AT5G22300 (*NIT4*), AT1G08980 (*AMI1*), AT3G17970 (*TOC64-III*), AT5G09420 (*TOC64-V*), AT5G64440 (*FAAH1*), AT5G07360 (*FAAH2*), AT3G25660 (*FAAH3*), AT4G34880 (*FAAH4*), AT2G38120 (*AUX1*), AT4G39950 (*CYP79B2*), AT2G22330 (*CYP79B3*), AT4G37550 (*IAMH1*), AT4G37560 (*IAMH2*).

## Acknowledgements

We are grateful to Drs. Paul Jarvis (Oxford University), Henrik Aronsson (University of Gothenburg), Kent Chapman (University of Northern Texas), Elison Blancaflor (Noble Foundation), and Hiroyuki Kasahara (RIKEN Center for Sustainable Resource Center, Japan) for kindly sharing mutant seeds with us. We also thank Dr. Jose Bruno-Barcena (NC State University, USA) for his assistance with plant sample lyophilization, Drs. Miguel Flores Vergara and Robert Franks for sharing reagents and equipment, and members of the Alonso-Stepanova laboratory and the INTRINSyC community at NCSU for their valuable feedback and discussions.

## Author contributions

A.N.S. and J.M.A. conceived the project. A.N.S., J.M.A., and M.F. designed the research. A.N.S., J.B., and M.F. generated Arabidopsis mutants. A.Pě. performed the quantitative profiling of auxin-related metabolites. B.E. performed quantitative analysis of gene expression of intronic T-DNA alleles. J.D. and A.Pa. contributed to the identification of high order mutants. X.L. tested the spontaneous conversion of IAOx into IAA *in vitro*. M.F. conducted experiments, analyzed the data, and prepared figures. S.B. contributed to the analysis of the results. A.N.S., J.M.A., K.L., N.O., and M.M.K. supervised research. M.F., A.N.S., and J.M.A. wrote the manuscript with the contributions of other authors.

---

**Supplementary Figure S1.**
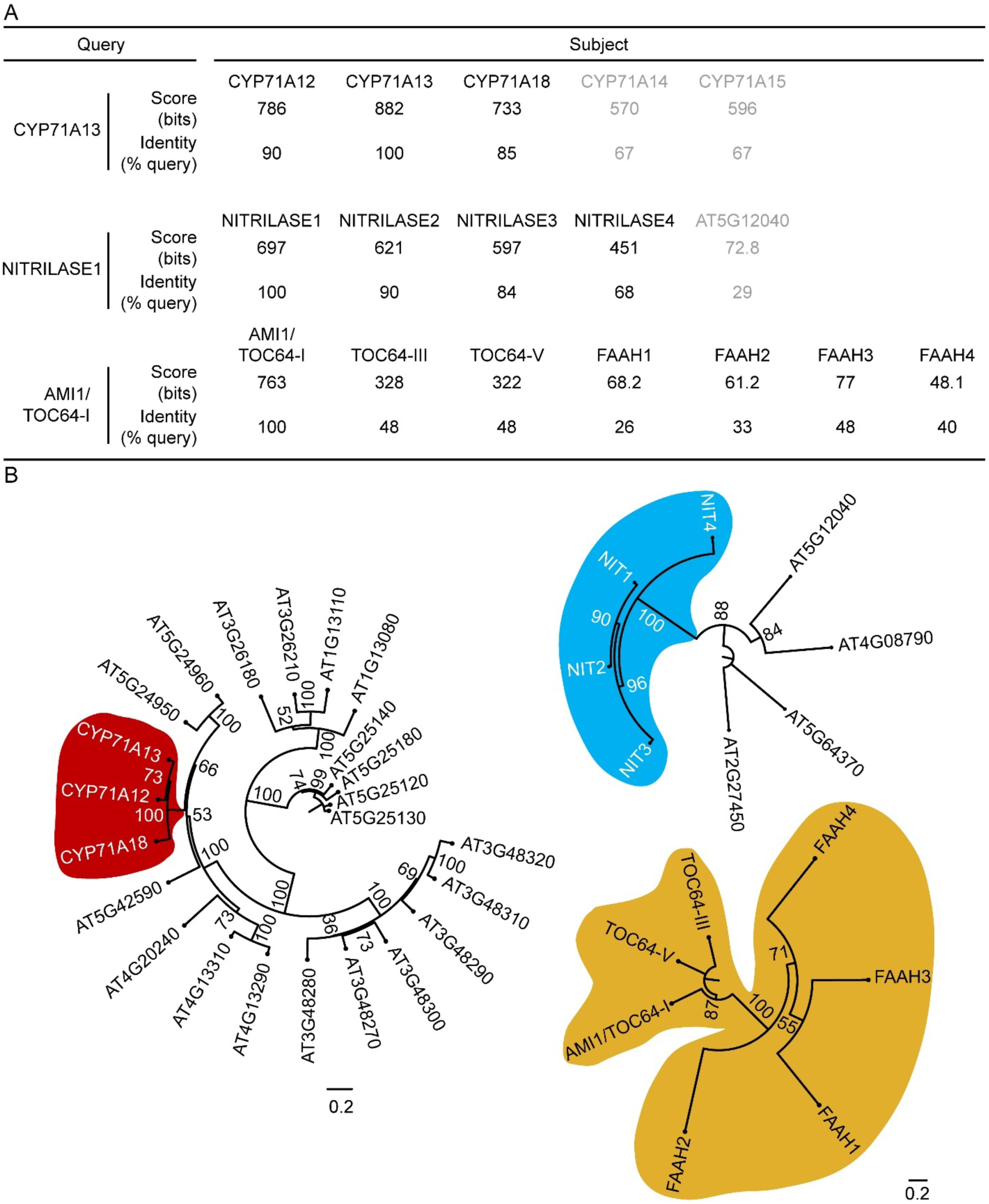
Phylogenetic comparison of amino acid sequences of Arabidopsis CYP71A, NIT, and AMI1/TOC64/FAAH family members. (A) Protein identity based on amino acid sequence comparison using protein basic local alignment search tool (BLASTP;(Camacho et al., 2009). Marked in gray are genes not considered for this study due to their lower identity score to the founding member of each family. (B) Maximum likelihood phylogenetic trees made using MUSCLE algorithm in PLAZA Dicots 5.0 (Van Bel et al., 2021). Node numbers represent bootstrap values.

**Supplementary Figure S2.**
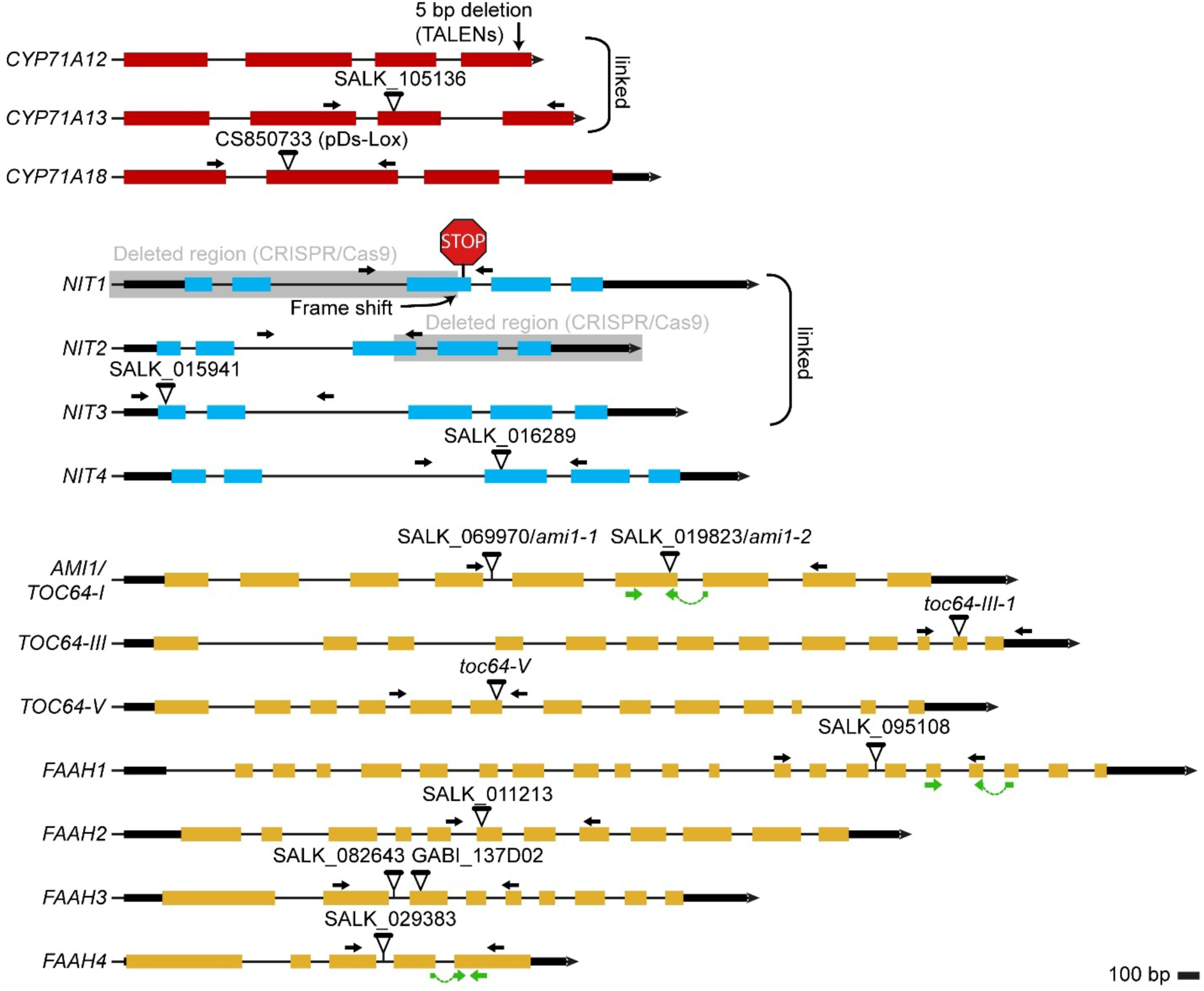
Schematic representation of gene structure (5’ to 3’) of *CYP71A*, *NIT*, and *AMI1/TOC64/FAAH* gene families examined in this study. Colored rectangles mark exons. Black rectangles denote 5’ and 3’ untranslated regions (UTRs). Black lines represent introns and the arrowhead at the right end of each gene model indicates the 5’-3’ orientation. T-DNA insertions (triangles) and primer annealing sites for genotyping (black arrows) and qPCR (green) are shown. Grey background in *NIT1* and *NIT2* corresponds to deleted DNA sequences in the CRISPR line.

**Supplementary Figure S3.**
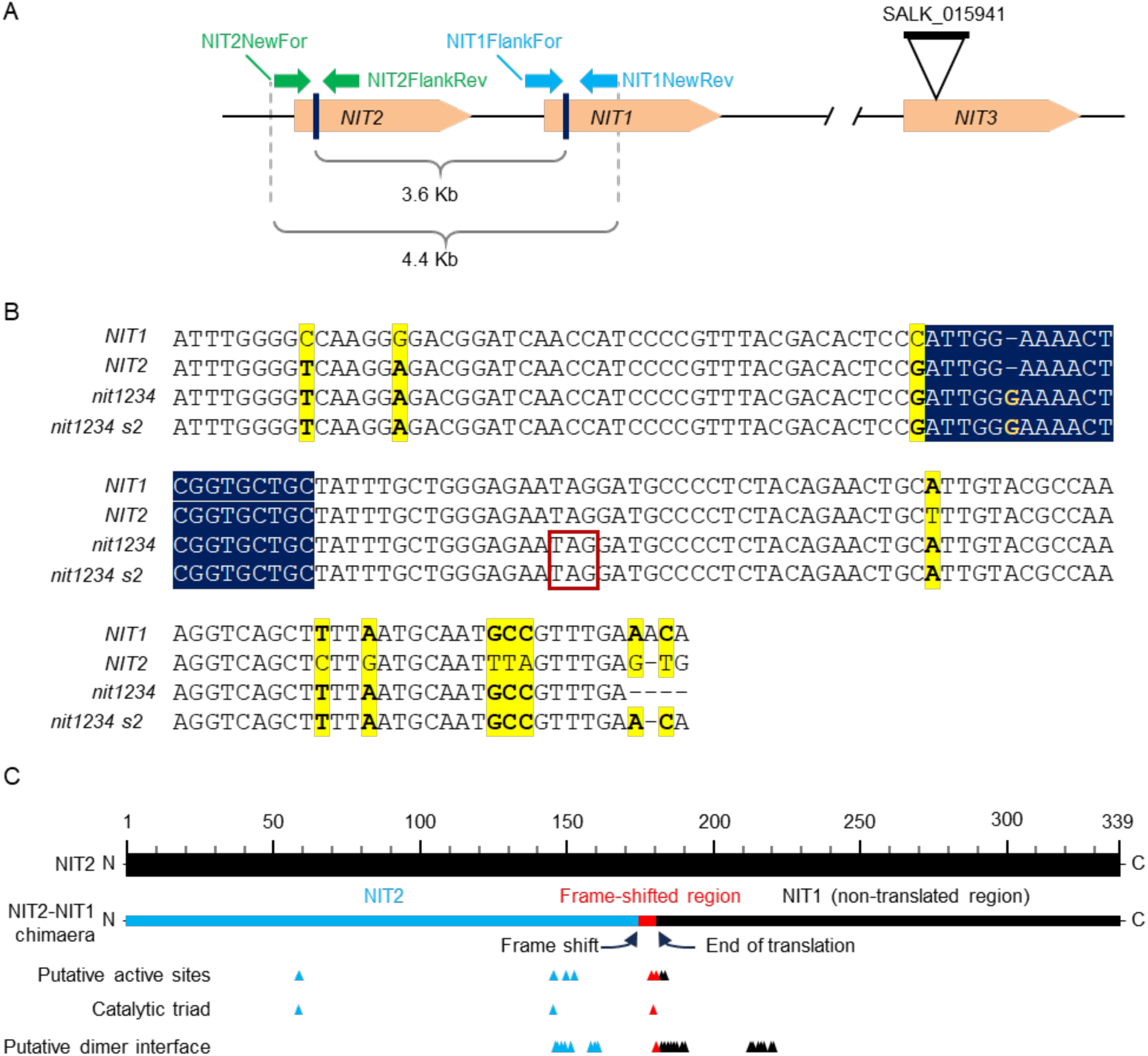
Genome editing of tandem *NIT2* and *NIT1* genes. (A) Relative positions of two genetically linked *NIT* genes, *NIT2* and *NIT1.* Distances between the two gRNA target sites (3.6 Kb), locations of PCR genotyping primers (colored arrows), and the size of the PCR fragment in WT background (4.4 Kb) are indicated. (B) Chimeric frameshifted *NIT2-NIT1* gene resulting from CRISPR/Cas9 genome editing. The gRNA targeting site (navy blue background) is the merging point between the two genes and contains a single G insertion that leads to a premature STOP codon (red rectangle). Polymorphisms between *NIT1* and *NIT2* (yellow background) show that upstream of the merging point, the PCR-amplified chimeric fragment contains *NIT2*-like polymorphisms, whereas downstream of the merging point, polymorphisms are *NIT1*-like, which suggests a complete deletion of the 3.6 Kb fragment depicted in panel A. An addition of a single G at the junction site shifts the reading frame and makes the chimeric gene non-functional. *NIT1* and *NIT2* sequences were obtained from TAIR. Chimeric sequences from *nit1234* and *nit1234 s2* were obtained by Sanger sequencing of the PCR fragment amplified using primers NIT2NewFor and NIT1FlankRev (primer sequences available in Supp. Table 3). *nit1234: nit1 nit2 nit3 nit4, s2: sur2.* (C) NIT2-NIT1 chimeric protein schematic showing the active sites predicted for NIT2 that are maintained (cyan) and lost (red and black) after the frame shift caused by the insertion of an extra G, as depicted in panel B. NIT2 amino acid residues that belong to putative active sites, catalytic triad, or putative dimer interface were obtained from the conserved domain database (Wang et al., 2023) accessed through BLASTP (Camacho et al., 2009) at the National Center for Biotechnology Information website (https://blast.ncbi.nlm.nih.gov/Blast.cgi).

**Supplementary Figure S4.**
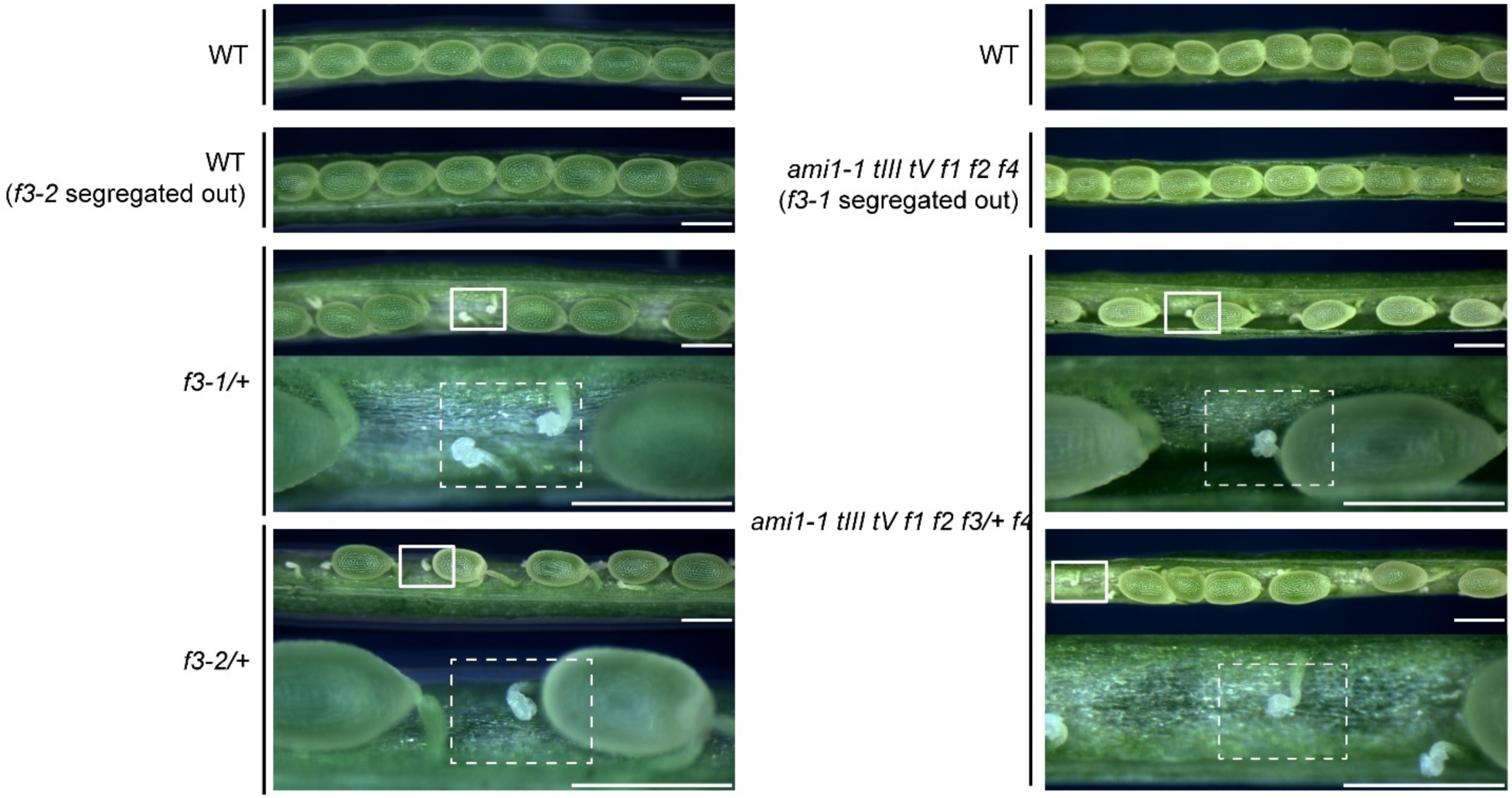
Embryo lethality in two independent *faah3* mutant alleles is observed in dissected siliques. The valves of fully expanded siliques (number 14-16 from the top) were stripped using Dumoxel Style 5 forceps under a Nikon SMZ645 dissection stereomicroscope and imaged using a QImaging MicroPublisher 5.0 RTV coupled to a Leica MZ125 dissection stereomicroscope. Scale bar=0.5mm. WT: wild-type (Col-0), *f3-1/+:* heterozygous *faah3-1* (SALK_082643), *f3-2/+*: heterozygous *faah3-2* (GABI_137D02), *ami1-1 tIII tV f1 f2 f4*: *ami1-1 toc64-III toc64-V faah1 faah2 faah4*, *ami1-1 tIII tV f1 f2 f3/+ f4: ami1-1 toc64-III toc64-V faah1 faah2 faah3-1/+ faah4.* Scale bar = 0.5mm.

**Supplementary Figure S5.**
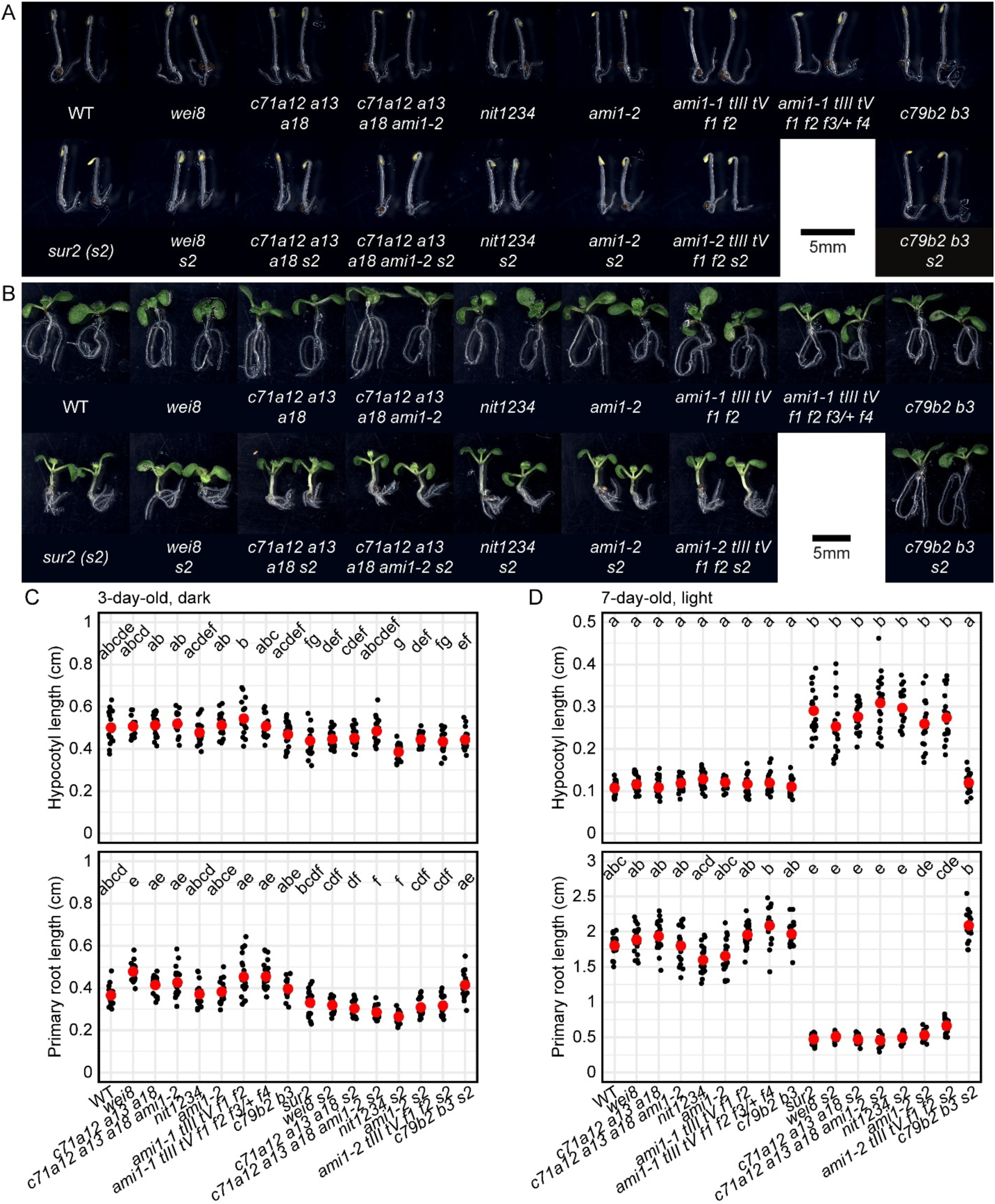
Mutants defective in the proposed IAOx pathway of auxin biosynthesis display no prominent growth defects. (A, B) Seedlings were germinated on horizontal plates in the dark for three days (A) or under continuous light for seven days (B). (C, D) Hypocotyl- and root-length quantification of seedlings grown for three days in the dark (C) or for seven days under continuous light (D). WT: wild-type (Col-0), *c71a12 a13 a18: cyp71a12 cyp71a13 cyp71a18, c71a12 a13 a18 ami1-2: cyp71a12 cyp71a13 cyp71a18 ami1-2, nit1234: nit1 nit2 nit3 nit4, ami1-1 tIII tV f1 f2: ami1-1 toc64-III toc64-V faah1 faah2, ami1-1 tIII tV f1 f2 f3/+ f4: ami1-1 toc64-III toc64-V faah1 faah2 faah3/+ faah4, s2: sur2, c79b2 b3: cyp79b2 cyp79b3*.

**Supplementary Figure S6.**
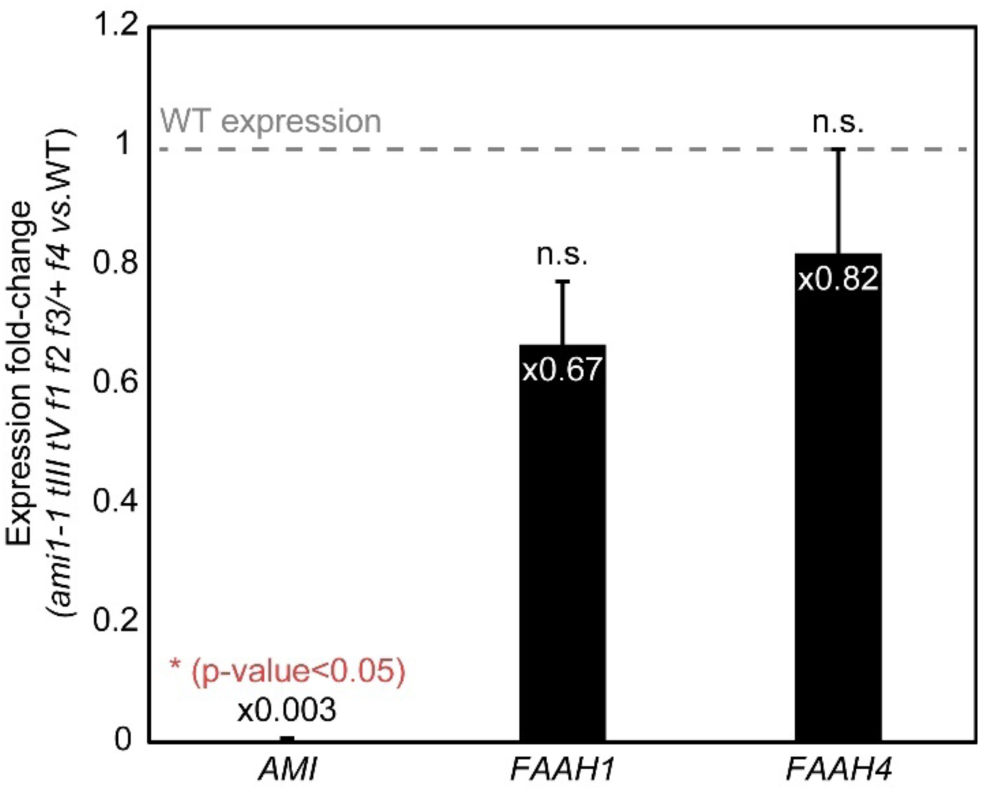
Expression analysis of intronic T-DNA alleles shows that *AMI1* is inactivated, but *FAAH1* and *FAAH4* are only partially knocked down. RNA was extracted from ten-day old WT and *ami1-1 toc64-III toc64-V faah1 faah2 faah3/+ faah4* (noted as *ami1-2 tIII tV f1 f2 f3/+ f4*) whole seedlings grown under continuous light on horizontal AT plates and analyzed by RT-qPCR. Statistical analysis (t-Student, α=0.05) was performed using ΔΔCт values, and expression fold-change was calculated as 2^(-ΔΔCт). ΔΔCт *GENE1* = ΔCт (*GENE1*_*mutant*) - ΔCт (*GENE1*_WT). ΔCт_genotype1 = Cт (*GENE1*_genotype1) - Cт (*CBP20*_genotype1)*. CBP20: CAP-BINDING PROTEIN20* (*AT5G44200*) used as a housekeeping gene to normalize gene expression.

**Supplementary Figure S7.**
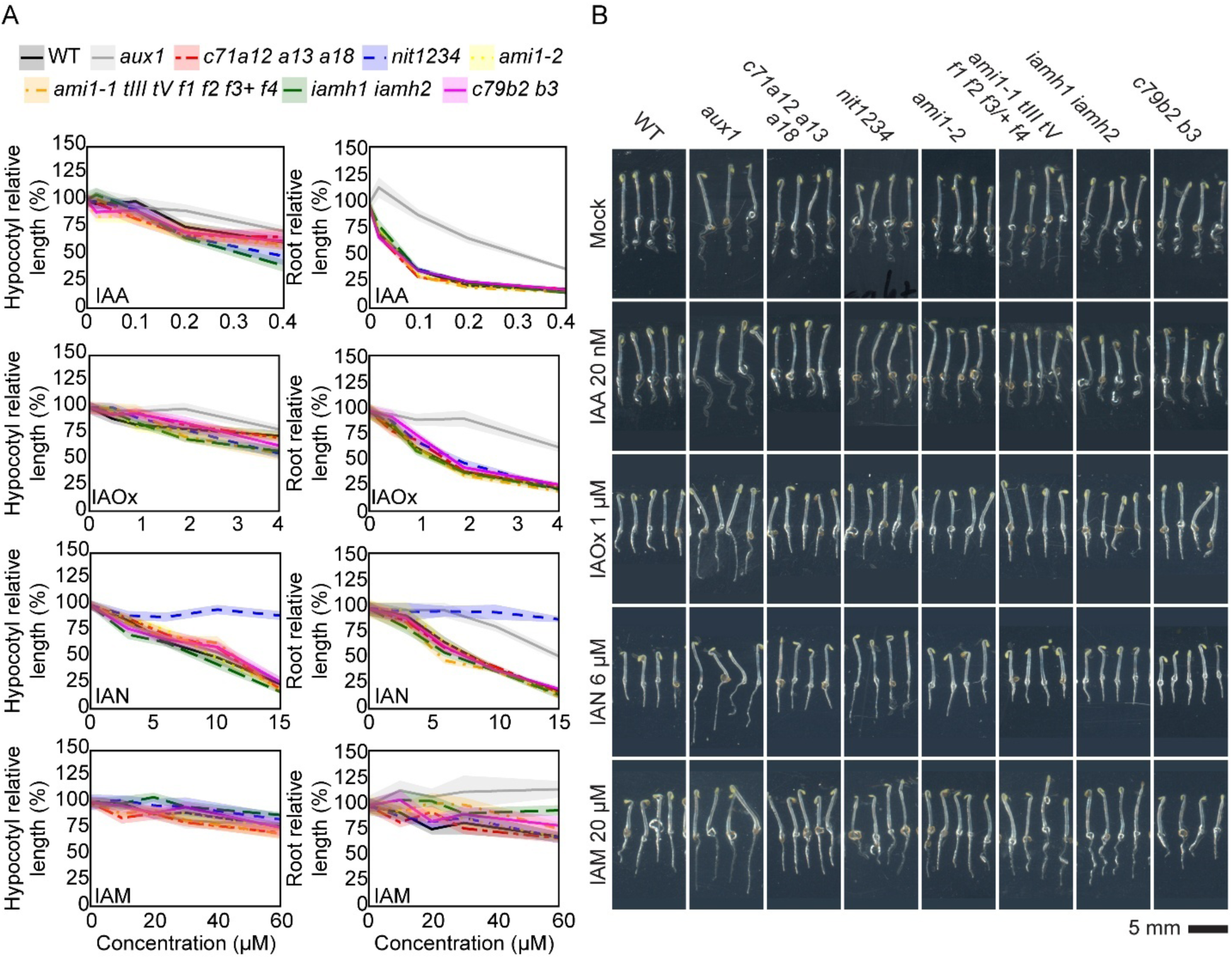
Phenotypes of dark-grown mutants impaired in the putative IAOx route challenge the established model of the IAOx pathway. (A) WT and mutant lines were germinated on horizontal plates in the dark for three days in control media and in media supplemented with the indicated concentrations of IAA, IAOx, IAN and IAM (Supp. Table 2). For each treatment, the “0” concentration contains the equivalent concentration of DMSO as the highest concentration tested for a specific precursor. Root and shoot lengths were measured in ImageJ. Relative organ size at a given concentration for a specific genotype was calculated by dividing the organ size by that in the corresponding control ([metabolite]=0). Average relative organ sizes (lines) and confidence intervals (CI=95%, shades) were plotted using R studio. (B) Photographs of representative plants for one of the concentrations for each compound. These precursor concentrations were chosen as they produce similar organ sizes in the WT as 20nM IAA. WT: wild-type (Col-0), *aux1: aux1-7, c71a12 a13 a18: cyp71a12 cyp71a13 cyp71a18, nit1234: nit1 nit2 nit3 nit4, ami1-1 tIII tV f1 f2 f3/+ f4: ami1-1 toc64-III toc64-V faah1 faah2 faah3/+ faah4, iamh1 iamh2: iamh1-1 iamh2-2, c79b2 b3: cyp79b2 cyp79b3*.

**Supplementary Figure S8.**
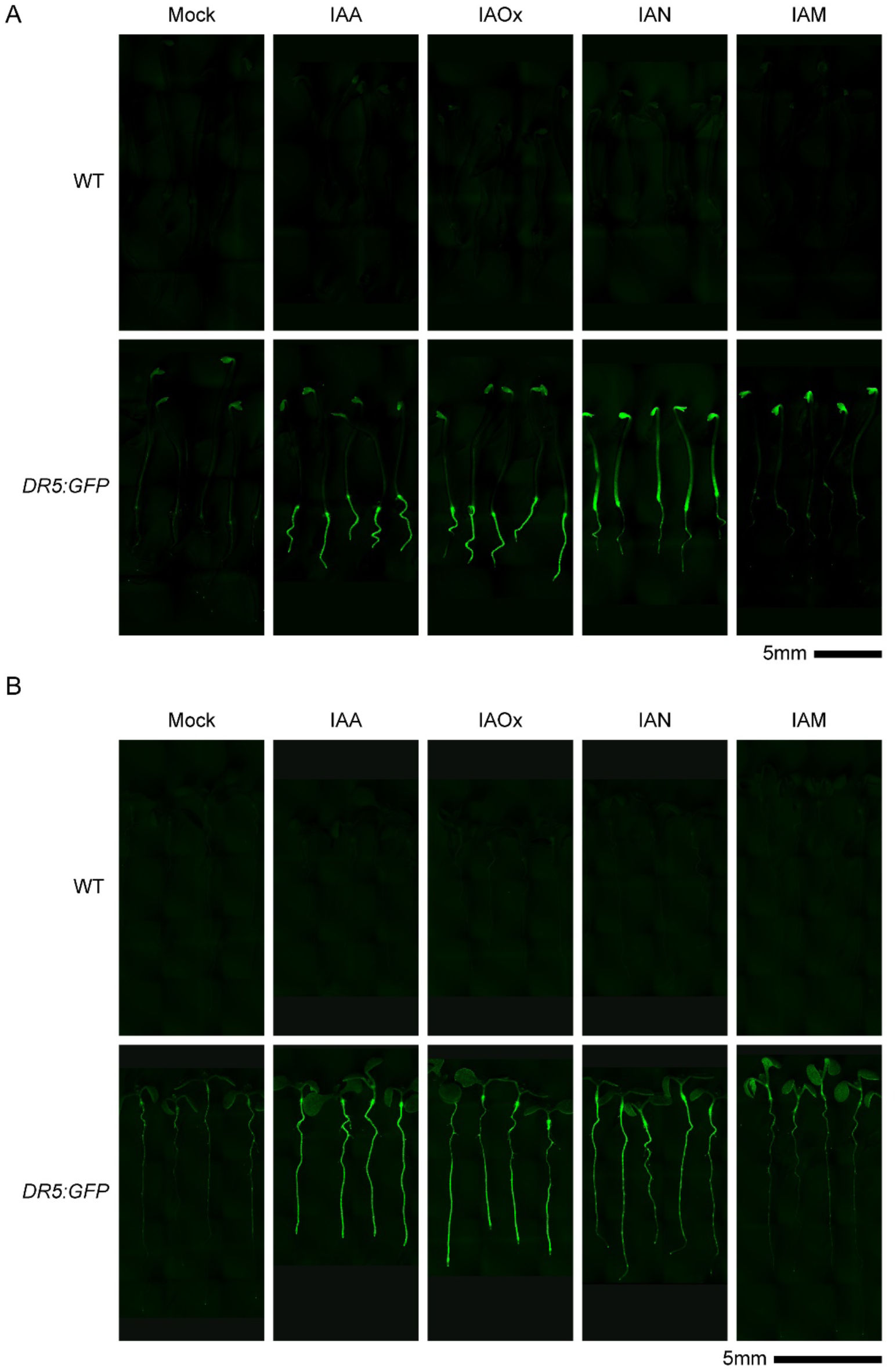
Exogenous application of putative IAOx intermediates induces activity of the auxin response reporter, *DR5:GFP*. (A,B) Three-day-old dark-grown seedlings (A) and five-day-old light-grown seedlings (B) were germinated on horizontal AT plates and transferred onto the indicated auxin precursors at concentrations empirically determined to be the lowest for seedlings to reach maximum change in organ size when germinated in the presence of that metabolite (Supplementary Fig. 6A). Dark experiment: 0.3µM IAA, 5µM IAOx, 20µM IAN, 50µM IAM, 0.25µM NAA. Light experiment: 3µM IAA, 20µM IAOx, 30µM IAN, 50µM IAM, 0.5µM NAA.

**Supplementary Figure S9.**
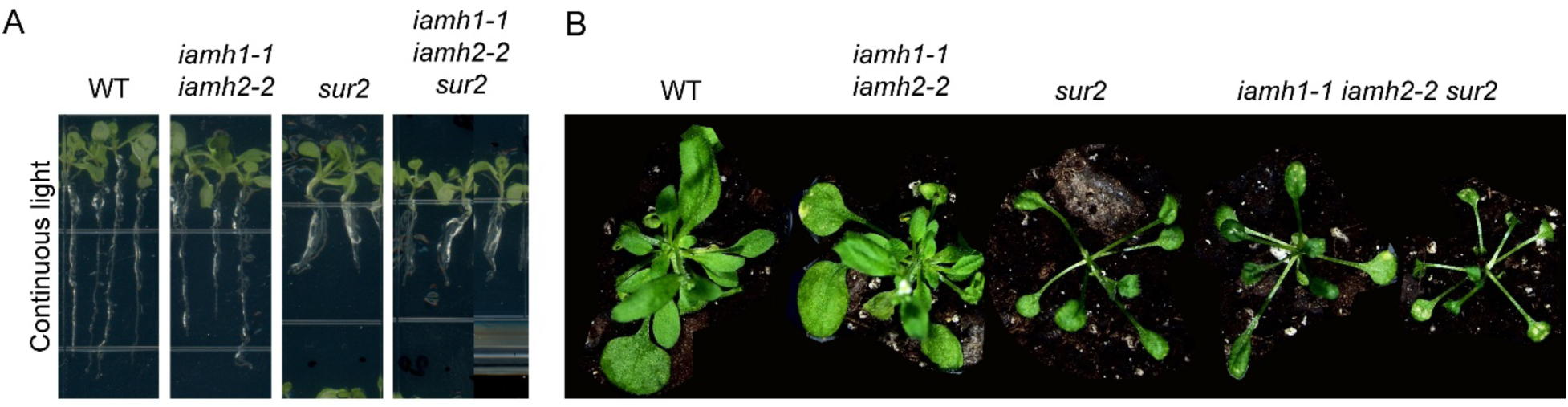
*iamh1 iamh2 sur2* mutants are indistinguishable from *sur2*. (A) Seeds from two *iamh1/+ iamh2/+ sur2* F2 plants (MF2842 and MF2845) were germinated and grown under continuous LED light for ten days on horizontal AT plates. *iamh1-1* and *iamh2-2* are genetically linked, and therefore, one fourth of the F2 progeny plants were *iamh1-1 iamh2-2 sur2*, as confirmed by genotyping. (B) Seedlings displayed in (A) were transferred to soil and grown for two additional weeks in long-day conditions (16-h light:8-h darkness) under white LED light prior to imaging. Plants that were genotyped as *iamh1 iamh2 sur2* triple mutants were confirmed by NcoI digestion for *iamh2-2* and by Sanger sequencing for *IAMH1* (Gao et al., 2020). Therefore, we concluded that *iamh1-1 iamh2-2* mutations fail to suppress *sur2* phenotype.

**Supplementary Figure S10.**
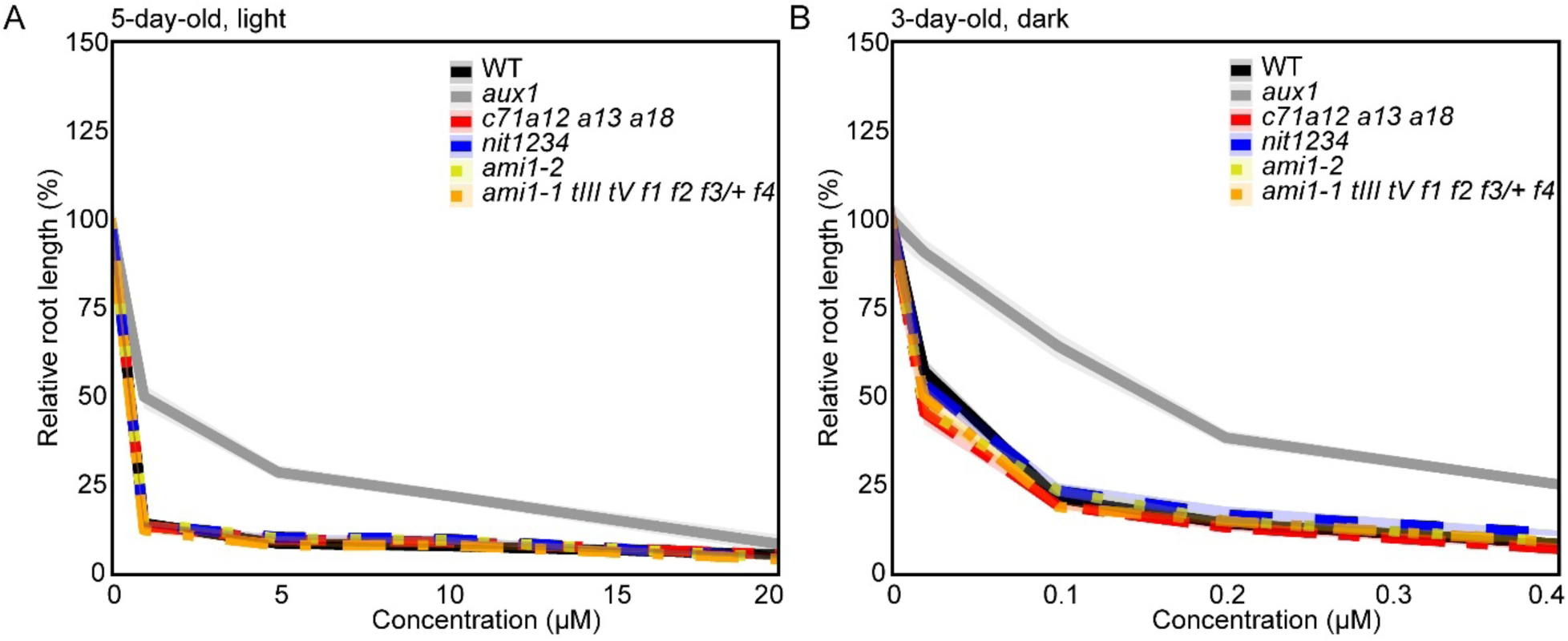
IAOx pathway mutants show normal root sensitivity to IAA. (A, B) An independent repetition of the experiments displayed in Figure 3 (A) and Supp. Figure 5 (B) showing that IAOx mutants possess normal root sensitivity to IAA both in the light and in the dark. WT: wild-type (Col-0), *c71a12a13a18: cyp71a12 cyp71a13 cyp71a18, nit1234: nit1 nit2 nit3 nit4, ami1-1 tIII tV f1 f2 f3/+ f4: ami1-1 toc64-III toc64-V faah1 faah2 faah3/+ faah4*.

**Supplementary Figure S11.**
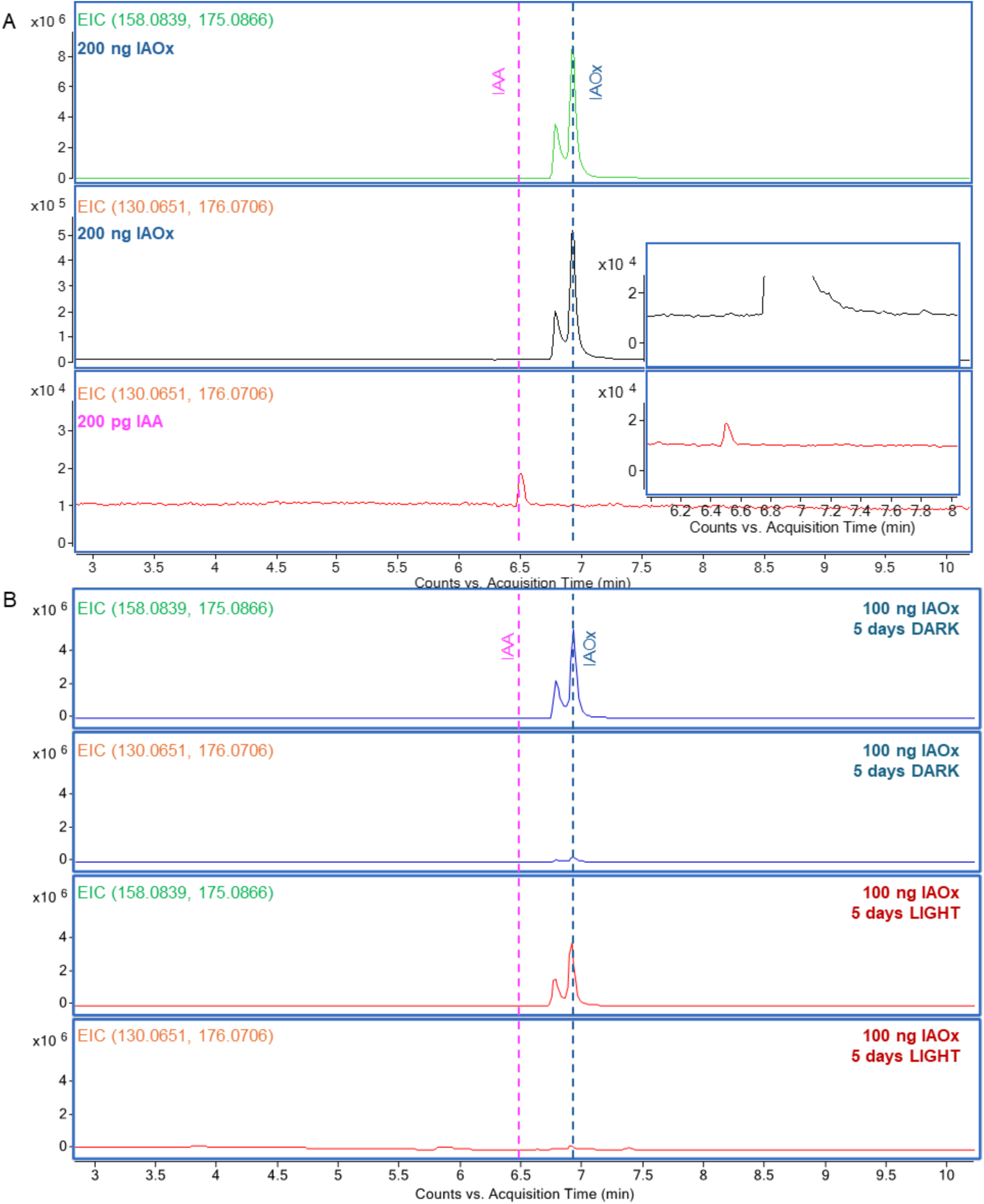
IAOx stock is not spontaneously converted into IAA. (A) Extracted ion chromatograms (EICs) of IAOx and IAA standards. EIC of 158.0839 + 175.0866 was used to detect IAOx; EIC of 130.0651 + 176.0706 was used to detect IAA. The insert shows the bottom two EICs at the same y axis scale. (B) An aqueous solution of IAOx (10 ng/uL) was incubated in a 22 °C growth chamber under light or in the dark for 5 days, and analyzed by LC-MS using the same method as in A.

**Supplementary Figure S12.**
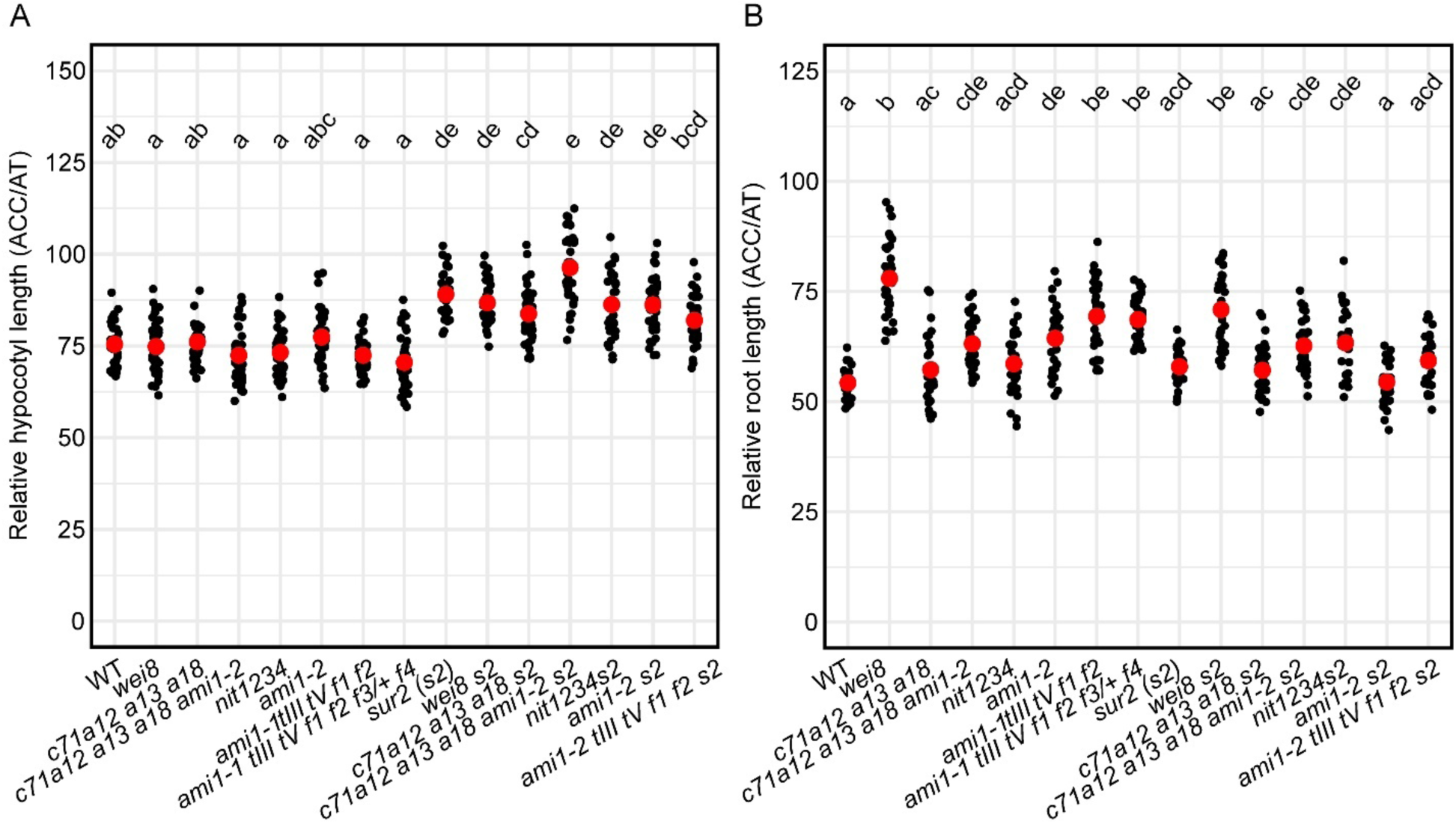
IAOx mutant hypocotyls and roots show WT-level of response to exogenous ACC. Examples of other experimental repetitions quantifying relative growth of hypocotyls (A) and roots (B) in response to ACC show that the mild defects observed in *cyp71a12a13a18s2* hypocotyls and *nit1234s2* roots in Figure5A-B are not consistently observed across repetitions and thus are unlikely to be biologically significant. Relative organ growth was calculated by dividing organ length of three-day-old, etiolated seedlings germinated on horizontal plates in the presence of the ethylene precursor ACC (0.2µM) by that in control media (AT). WT: wild-type (Col-0), *c71a12 a13 a18: cyp71a12 cyp71a13 cyp71a18, c71a12 a13 a18 ami1-2: cyp71a12 cyp71a13 cyp71a18 ami1-2, nit1234: nit1 nit2 nit3 nit4, ami1-1 tIII tV f1 f2: ami1-1 toc64-III toc64-V faah1 faah2, ami1-1 tIII tV f1 f2 f3/+ f4: ami1-1 toc64-III toc64-V faah1 faah2 faah3/+ faah4, s2: sur2*.

**Supplementary Figure S13.**
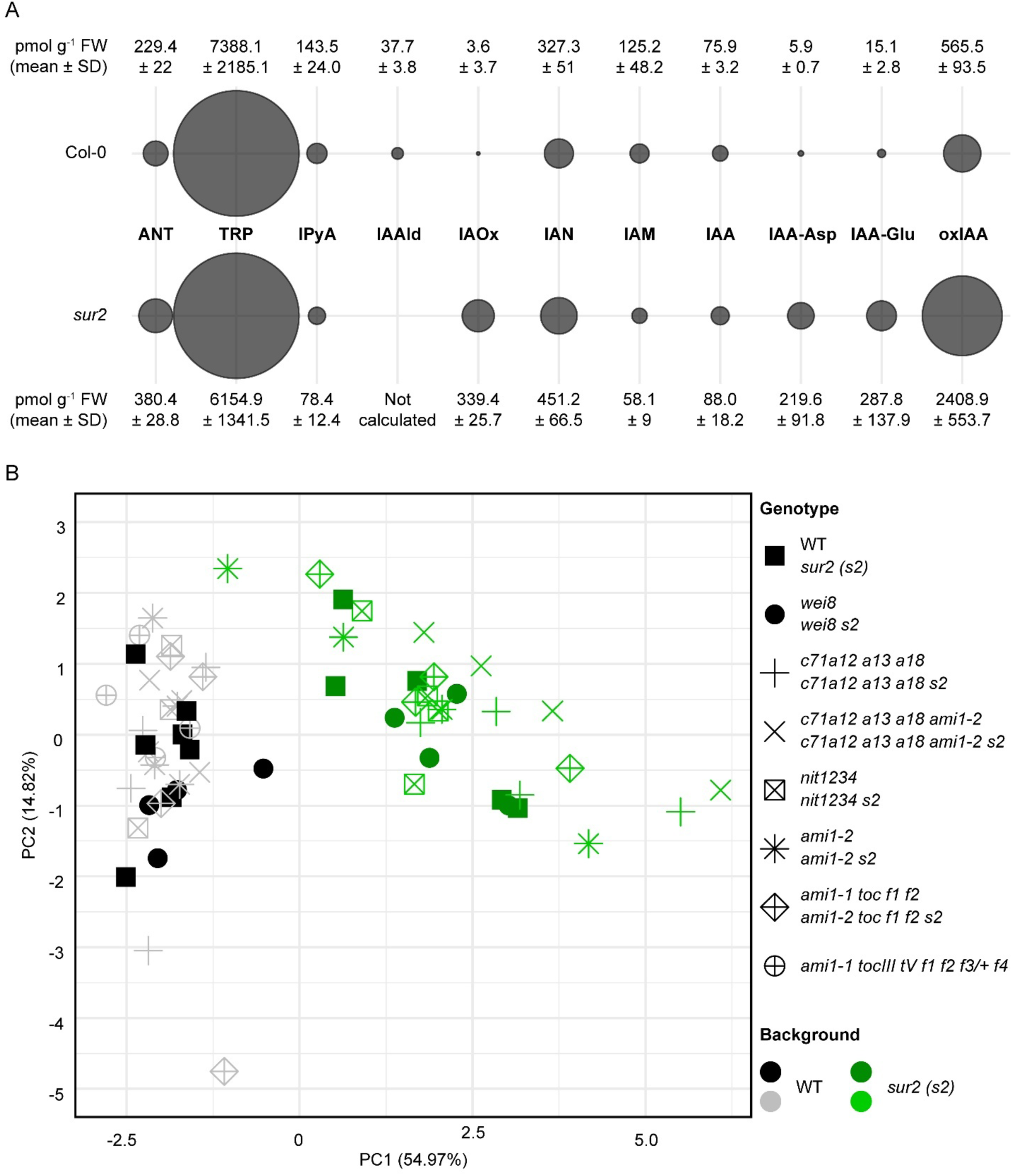
*sur2* background is the only mutation tested that prominently impacts the Arabidopsis metabolic profiles in seedlings. (A) Absolute values of metabolites measured in pmol/g FW of WT (Col-0) and *sur2*. (B) Principal Component Analysis of metabolic profiles for all metabolites (not including IAAld or TAM) in the mutants. <LOD values were substituted by the average value of the other replicates when significantly different from 0 (for example, for IAOx <LOD are substituted by 0). WT: wild-type (Col-0), *c71a12 a13 a18: cyp71a12 cyp71a13 cyp71a18, c71a12 a13 a18 ami1-2: cyp71a12 cyp71a13 cyp71a18 ami1-2, nit1234: nit1 nit2 nit3 nit4, ami1-1 tIII tV f1 f2: ami1-1 toc64-III toc64-V faah1 faah2, ami1-1 tIII tV f1 f2 f3/+ f4: ami1-1 toc64-III toc64-V faah1 faah2 faah3/+ faah4, s2: sur2*.

**Supplementary Figure S14.**
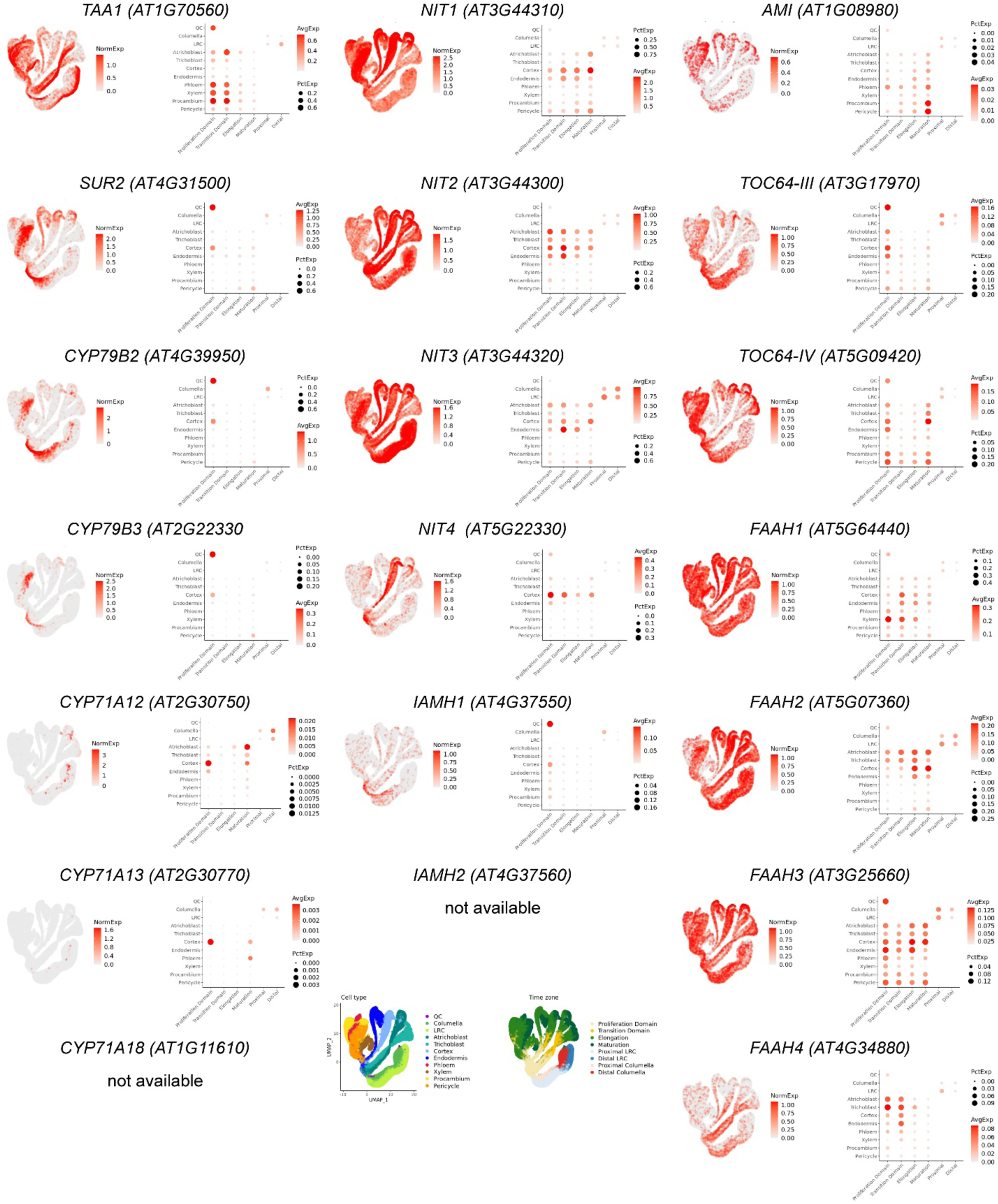
Single-cell gene expression analysis shows an overlap in the cortex cells, but not in the quiescent center (QC), between the gene families studied in this work. Cells were collected from 5-day-old Arabidopsis WT roots from seedlings grown under long-day conditions. Data obtained from Nolan et al., 2023 and visualized using *Arabidopsis* Root Virtual Expression eXplorer (ARVEX; https://shiny.mdc-berlin.de/ARVEX/). NormExp: Normalized gene expression. AvgExp: Averaged normalized gene expression value in specific time zone & cell type. PctExp: Percentage of cells that the gene is expressed in specific time zone & cell type. Quantile cutoff (0.1-0.9): minimum and maximum cutoff values specified as quantiles for non-zero normalized gene expression. LRC: lateral root cap.

