## Supplementary tables for "The *CYP71A*, *NIT*, *AMI,* and *IAMH* gene families are dispensable for indole-3-acetaldoxime-mediated auxin biosynthesis in Arabidopsis"

**Supplementary Table 1**. Mutant lines used or generated in this work and their stock numbers.

| Mutant line | Source |
|---|---|
| *wei8-1* | CS16407; Stepanova et al., 2008 |
| *cyp71a12 cyp71a13 cyp71a18* | This work |
| *cyp71a12 cyp71a13 cyp71a18 ami1-2* | This work |
| *nit1 nit2 nit3 nit4* | This work |
| *ami1-2* | SALK_019823; Pérez-Alonso et al., 2020 |
| *ami1-1 toc64-III toc64-V faah1 faah2* | This work |
| *ami1-1 toc64-III toc64-V faah1 faah2 faah3/+ faah4* | This work |
| *sur2 DR5:GUS* | CS16401; Stepanova et al., 2005 |
| *wei8-1 sur2* | CS16437; generated in Stepanova et al., 2008, characterized in this work |
| *cyp71a12 cyp71a13 cyp71a18 sur2* | This work |
| *cyp71a12 cyp71a13 cyp71a18 ami1-2 sur2* | This work |
| *nit1 nit2 nit3 nit4 sur2* | This work |
| *ami1-2 sur2* | This work |
| *ami1-2 toc64-III toc64-V faah1 faah2 sur2* | This work |

**Supplementary Table 2**. Primer combinations used for genotyping and PCR product size.

| Allele | Mutant line | Mutant allele | WT allele |
|---|---|---|---|
| *wei8-1* | Act tag T-DNA<br>(Stepanova et al., 2008) | DWLB1+7056-R1<br>~800 bp | 70560-F8+70560-R1<br>850 bp |
| *aux1-7* | 3218G>A<br>(Pickett et al., 1990) | AUX1_F+aux1_EcoRV_R, then digest with EcoRV<br>330 bp | 355 bp |
| *cyp79b2* | Pooled SALK collection<br>(Zhao et al., 2002) | 79b2-F+JMLBa<br>>803 bp | 79b2-F+79b2-R<br>803 bp |
| *cyp79b3* | Pooled SALK collection<br>(Zhao et al., 2002) | JMLBa+79b3-R<br>>378 bp | 79b3-F+79b3-R<br>378 bp |
| *sur2* | SALK_028573<br>(Alonso et al., 2003; Stepanova et al., 2005) | CYP83B1-F1+JMLBa<br>~1000 bp | CYP83B1-F1+CYP83B1-R1<br>803 bp |
| *cyp71a12* | *cyp71a12*$^{TALEN}$*/a13n*<br>(Müller et al., 2015) | A 5 bp deletion genetically linked to *cyp71a13* | |
| *cyp71a13* | SALK_105136<br>(this work; Alonso et al., 2003) | JMLB1+71A13-R2a<br>>839 bp | JMLB1+71A13-F2a+71A13-R2a<br>1163 bp |
| *cyp71a18* | WiscDsLox297300_18A<br>(this work; Woody et al., 2007) | WiscLB1+71A18-R1<br>>341 bp | MF_71A18-F2+71A18-R1<br>778 bp |
| *nit1* | *NIT2/NIT1*$^{CRISPR}$<br>(this work) | NIT2NewFor + NIT1NewRev<br>692 bp | NIT1FlankFor+NIT1NewRev<br>586 bp |
| *nit2* | | | NIT2NewFor+NIT2FlankRev<br>669 bp |
| *nit3* | SALK_015941<br>(this work; Alonso et al., 2003) | NIT3-F2+JMLBa<br>>254 bp | NIT3-F2+NIT3-R2<br>801 bp |
| *nit4* | | NIT4-F1+JMLBa | NIT4-F1+NIT4-R1 |

| | | | |
|---|---|---|---|
| | SALK_016289 (this work; Alonso et al., 2003) | >501 bp | 832 bp |
| *ami1-1/ami1-2* | *ami1-1*: SALK_069970 (Alonso et al., 2003; Aronsson et al., 2007; Pérez-Alonso et a., 2020) | JMLB1+AMI1-R2 | AMI1-F2+AMI1-R2 |
| | *ami1-2*: SALK_019823 (Alonso et al., 2003; Pérez-Alonso et a., 2020) | >838 bp for *ami1-1* and >400 bp for *ami1-2* | 916 bp |
| *toc64-III* | *toc64-III-1* (Rios et al., 2002; Aronson et al., 2007) | mut 64-III-LB+64-III-R(PJ) | 64-III-F+64-III-R(PJ) |
| | | >700 bp | 642 bp |
| *toc64-V* | Garlic_565_D12 (Session et al., 2002; Aronson et al., 2007) | SAIL-LB3+64-V-R | 64-V-F+64-V-R |
| | | >200 bp | 588 bp |
| *faah1* | SALK_095108 (Alonso et al., 2003; Wang et al., 2008; Keereetaweep et al, 2013) | FAAH1-F+JMLBa | FAAH1-F+FAAH1-R |
| | | >500 bp | 887 bp |
| *faah2* | SALK_011213 (Alonso et al., 2003; Keereetaweep et al, 2013) | JMLB1+FAAH2-R | FAAH2-F+FAAH2-R |
| | | >520 bp | 652 bp |
| *faah3-1* | SALK_082643 (this work; Alonso et al., 2003) | FAAH3-F+JMLB1 | |
| | | >450 bp | FAAH3-F+FAAH3-R |
| *faah3-2* | GABI_137D02 (this work; Rosso et al., 2003) | DWLB1+FAAH3-R | 852 bp |
| | | >450 bp | |
| *faah4* | SALK_029383 (this work; Alonso et al., 2003) | JMLBa+FAAH4-R2 | FAAH4-F2+FAAH4-R2 |
| | | >700 bp | 690 bp |
| *iamh1* | *iamh1-1 iamh2-2*$^{CRISPR}$ (Gao et al., 2020) | iamh1-1_F+iamh1-1_R, then digest with NcoI | |
| | | 1500 bp | 600bp+900 bp |
| *iamh2* | *iamh1-1 iamh2-2*$^{CRISPR}$ (Gao et al., 2020) | SNP genetically linked to *iamh1-1*. DNA sequencing. | |

**Supplementary Table 3**. Primer sequences used for mutant genotyping.

| Target | Name | DNA sequence |
|---|---|---|
| T-DNA LB | JMLB1 | GGCAATCAGCTGTTGCCCGTCTCACTGGTG |
| | JMLBa | CTTTGACGTTGGAGTCCACGTTC |
| | SAIL-LB3 | TAGCATCTGAATTTCATAACCAATCTCGATACAC |
| | WISC-LB1 | AACGTCCGCAATGTGTTATTAAGTTGTC |
| | TOC64-III-1 LB | GTTGACAGACTGCCTAGCATTTGAGTG |
| | DWLB1 | CATACTCATTGCTGATCCATGTAGATTTCC |
| *TAA1* | 70560-F8 | CATCAGAGAGACGGTGGTGAAC |
| | 70560-R1 | GCTTTTAATGAGCTTCATGTTGG |
| *AUX1* | AUX1_F | GAGAGTCTGAGTATGACAAATC |
| | aux1_EcoRV_R | TACATTGGTAACACTTGGCAAAGATA |
| *C79B2* | C79B2-F | AGTATCATGACCCAATCATCGAC |

| | | |
|---|---|---|
| | C79B2-R | CCATATCGGCTAAGAAGGAC |
| *C79B3* | C79B3-F | GCAATCCACCAATATCCGTCAG |
| | C79B3-R | GTTCTATGCATGGACTCGTGG |
| *SUR2* | CYP83B1-F1 | GAGACTCTTGACCCTAACCGC |
| | CYP83B1-R1 | GCGAGTCCAGTCATGACGTCC |
| *C71A13* | 71A13-F2a | ATGGATAGATGGGATCCGTGG |
| | 71A13-R2a | GGCAAACATCGATACCAATGGC |
| *C71A18* | MF_71A18-F2 | GATGAAGGTATTTCATTACTAAGCCC |
| | 71A18-R1 | GACGAAATCCGCTTTATGTTCGCC |
| *NIT1* | NIT1FlankFor | GGCTATGGTTGGAGTCTTGG |
| | NIT1NewRev | AGCCTAGATGTTTCAAACGGC |
| *NIT2* | NIT2NewFor | AGGTTTGGACACTGATCCGT |
| | NIT2FlankRev | TTGCATCAAGAGCTGACCTTT |
| *NIT3* | NIT3-F2 | ATCCACCACCGGTTCGTTCTGC |
| | NIT3-R2 | GATCCGATTCTAACGGATCCTG |
| *NIT4* | NIT4-F1 | TTGGTTGCTCCACCCGTGAC |
| | NIT4-R1 | GAAGGATAGTCTTTCCGACGAC |
| *AMI1* | AMI1-F2 | TCCAATGGCTCAGAGCTTCG |
| | AMI1-R2 | CCACATTAGCTTGGAGATGCG |
| *TOC64-III* | TOC64-III-F | CCAAAGCCATCACCCTCGAC |
| | TOC64-III-R(PJ) | GACATGACCTTATTTTTGACGCTACGCTGAC |
| *TOC64-V* | TOC64-V-F | CATGCTACTCTAGGTGTTTGCC |
| | TOC64-V-R | GTAACCTCCAGCAAAGAGGG |
| *FAAH1* | FAAH1-F | GCAATGCAATAGGATCTCTACGAC |
| | FAAH1-R | GAGGTATCACTGGAGCTGTC |
| *FAAH2* | FAAH2-F | CTTGGTATCTTTCTCTAGTGCCG |
| | FAAH2-R | GGTATGTCATTGAGCCTGCTG |
| *FAAH3* | FAAH3-F | GCTGTTGCAGCAAGGCAGTG |
| | FAAH3-R | CTCCAAATCCTTCTCCACGG |
| *FAAH4* | FAAH4-F2 | CTGCGTCCGAGTTCATACCTG |
| | FAAH4-R2 | GGATACCCTCCAATGGCTAGG |
| *IAMH1* | iamh1-1_F | GATGACGCCAAGCGTGTAAGC |
| | iamh1-1_R | CTGGGAATTCAGAGGTAAGCAC |
| *IAMH2* | iamh2_F | GCATATAGGTAAAAGTTTCTTGGTGATTA |
| | iamh2_R | TCTGCTTGACAAAAAACAGGATGA |

**Supplementary Table 4**. Concentrations (µM) of auxin biosynthesis precursors used in assays shown in Figure 3 (A) and Supp. Figure 6 (B).

A) Continuous light (5 days)

| | | | | | |
|---|---|---|---|---|---|
| IAOx | 40 | 20 | 10 | 5 | Mock (DMSO like in 40 µM) |
| IAN | 30 | 25 | 20 | 15 | Mock (DMSO like in 30 µM) |

| | | | | | |
|---|---|---|---|---|---|
| IAM | 60 | 30 | 20 | 10 | Mock (DMSO like in 60 µM) |
| IAA | 20 | 10 | 5 | 1 | Mock (DMSO like in 20 µM) |

B) Continuous darkness (3 days)

| | | | | | |
|---|---|---|---|---|---|
| IAOx | 4 | 2 | 1 | 0.5 | Mock (DMSO like in 4 µM) |
| IAN | 15 | 10 | 6 | 3 | Mock (DMSO like in 15 µM) |
| IAM | 60 | 30 | 20 | 10 | Mock (DMSO like in 60 µM) |
| IAA | 0.4 | 0.2 | 0.1 | 0.02 | Mock (DMSO like in 0.4 µM) |

**Supplementary Table 5**. Primer combinations used for expression analysis of intronic T-DNA alleles by RT-qPCR.

| Target | Name | DNA sequence | Amplicon size |
|---|---|---|---|
| *AMI1/ami1-1* (SALK_069970) | ami1-1_flk_F | GCTTCGACACAGTTGGATGGT | 249 bp |
| | ami1-1_flk_R | CATTCTGTCCAATGTACTCTCCAA | |
| *FAAH1/faah1* (SALK_095108) | faah1_flk_F | CAATCACGGTTGCAAAGTGGTG | 164 bp |
| | faah1_flk_R | GCAAAGCTGGTACGAGTGTC | |
| *FAAH4/faah4* (SALK_029383) | f4_flk_F | CAGGACAGTATTGGGCCGAT | 245 bp |
| | f4_flk_R | ATGACAATAGCGCCTTCTCG | |
| *CBP20* (*AT5G44200*) | *CBP20*_F | AATCGCCATGGAAGAGGAGAC | 144 bp |
| | *CBP20*_R | GAATCGTGGGTTCTTCTCCGGTC | |

The *faah3-1* intronic mutant allele was not included in this analysis as it is embryo-lethal in homozygotes and phenotypically similar to the *faah3-2* exonic allele (Supplementary Figure 4).
